# Glucocorticoid receptor activation during human microglial differentiation leads to genomic instability and senescence

**DOI:** 10.1101/2022.05.23.493044

**Authors:** Jingzhang Wei, Charles Arber, Selina Wray, John Hardy, Thomas M Piers, Jennifer M Pocock

## Abstract

Early life stress, prenatal exposure to glucocorticoids (GCs), confers a higher risk of psychiatric and neurodevelopmental disorders in children. Increasingly, the importance of microglia in these disorders has been recognised. Studies on GCs exposure during microglial development have been limited, and there are few, if any, human studies. We established an *in vitro* model of ELS by continuous pre-expoure of human iPS-microglia to GCs during primitive haematopoiesis (the critical stage of iPS-microglial differentiation) and then examined how this exposure affected the microglial phenotype as they differentiated and matured to microglia. The iPS-microglia predominately expressed glucocorticoid receptors over mineralocorticoid receptors, and the GR-α splice variant. Chronic GCs exposure during primitive haematopoiesis was able to recapitulate *in vivo* ELS effects. Thus pre-exposure to prolonged GCs resulted in increased type I interferon signalling, the presence of Cyclic GMP-AMP synthase-positive (cGAS) micronuclei, and cellular senescence in the matured iPS-microglia. The findings from this *in vitro* ELS model have ramifications for the responses of microglia in the pathogenesis of GC-mediated ELS-associated disorders such as schizophrenia, attention-deficit hyperactivity disorder and autistic spectrum disorder.

**Highlights:**

- Human iPS-derived-microglia predominantly express glucocorticoid receptor NR3C1 compared with mineralocorticoid receptor NR3C2, and a predominant splice variant of the NR3C1 of GR-α.
- GC expression shows a differentiation-linked increment from iPSC to iPS-microglia.
- An early-life stress model was established by exposing iPSC to glucocorticoids during primitive haematopoiesis.
- RNA-seq analysis revealed that this early glucocorticoid exposure led to enhanced type I interferon inducible gene expression in the subsequent iPS-microglia.
- Furthermore, micronuclei formation and cellular senescence markers were upregulated in the iPSC-microglia, indicating genomic instability due to early chronic GC exposure.
- These findings have ramifications for the microglial responses in ELS linked neurodevelopmental disorders such as schizophrenia, attention-deficit hyperactivity disorder and autistic spectrum disorder.

**Graphical abstract:** 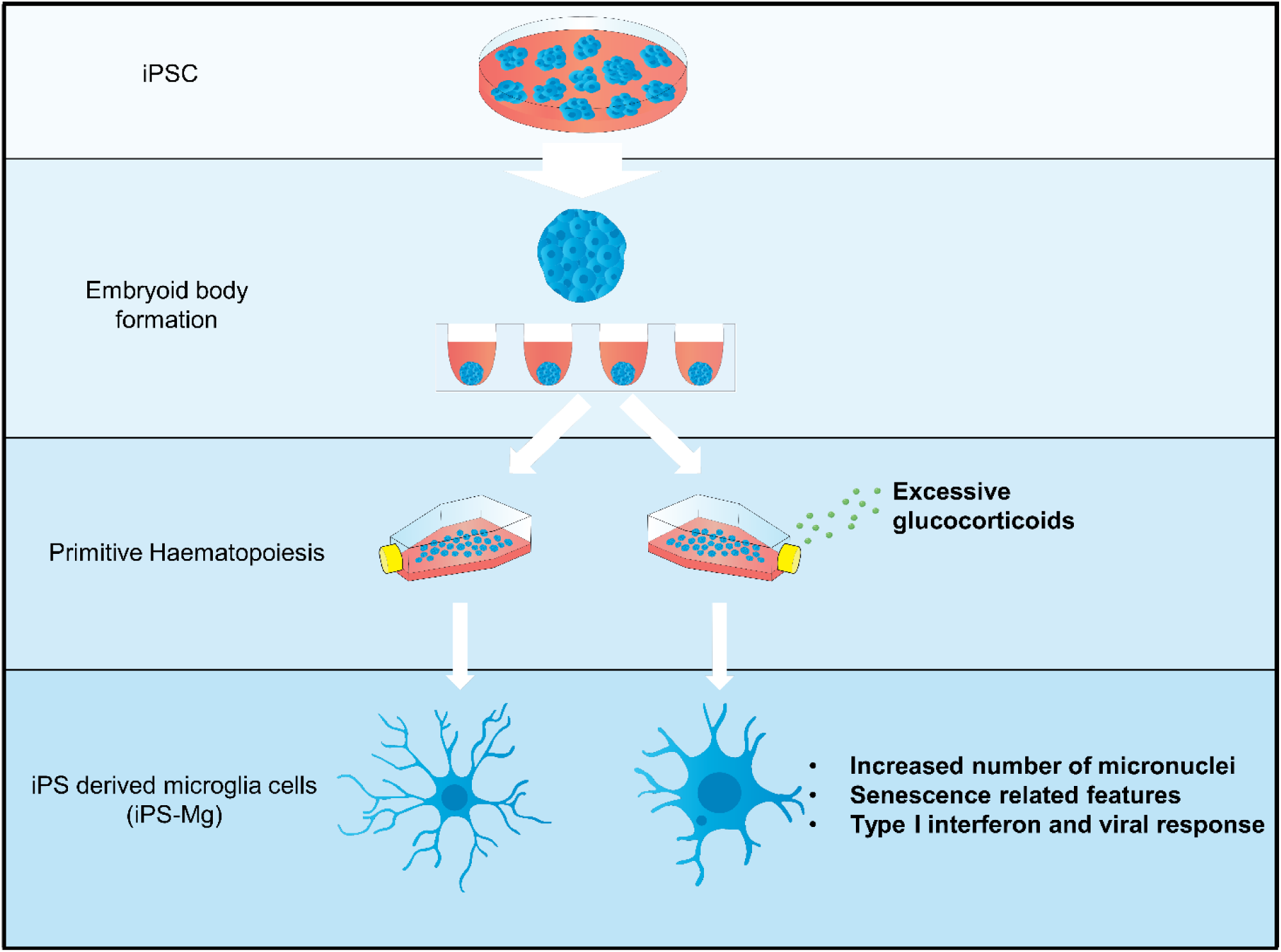

## Introduction

Early life stress (ELS) is defined as an adverse environment or experience that an individual encounters in critical developmental periods. ELS, such as prenatal/maternal stress, has been significantly associated with increased risk of neurodevelopmental and psychiatric disorders later in life^1, 2^ (Lupien et al. 2009, Manzari et al. 2019). One molecular mediator of ELS is prolonged elevated maternal glucocorticoids (GCs) during pregnancy and early life development^3^ (Krontira et al. 2020), and thus *in utero* chronic activation of its cognate receptors, glucocorticoid receptors (GR). There is a well-established relationship between dysregulation of maternal GCs/GR signalling and increased GC exposure to the foetal system^4,5,6,7^ (Benediktsson et al. 1993, Swanson and David 2015, Quinn et al. 2019, Chen et al. 2021).

Microglia are the primary immunocompetent cells within the central nervous system (CNS). During pre- and post-natal neurodevelopment, microglia participate in many neurodevelopmental processes, such as cortical neurogenesis^8,9,10,11^ (Schafer et al. 2012, Cunningham et al. 2013, Squarzoni et al. 2014, Weinhard et al. 2018), controlling migration of inhibitory neocortical interneurons that regulate excitation-inhibition balance^10^ (Squarzoni et al. 2014), and remodelling of synaptic structures^8, 11^ (Schafer et al. 2012, Weinhard et al. 2018). Mounting evidence points to the involvement of microglia in psychiatric and neurodevelopmental disorders, including, schizophrenia^12^ (Mondelli et al. 2017), autism spectrum disorder and attention-deficit hyperactivity disorder (ADHD)^13^ (Courchesne et al. 2020).

Microglial development occurs in early embryonic life. Microglia are derived from the embryonic yolk sac, and first appear in the CNS around gestational week 4. 5 in humans^14^ (Monier et al. 2007), and embryonic day 9. 5 in mice^15^ (Ginhoux et al. 2010). Therefore, we hypothesise that increased GC levels caused by prenatal GC overexposure could influence early microglial development and subsequent microglial phenotypes. In rat models, prenatal elevated GC was found to increase the overall density and immunoreactivity of embryonic microglia^16^ (Bittle and Stevens 2018). Furthermore, aberrant GR activation, the cognate GC receptor, in early postnatal primary microglia reduces CX3CR1 and TREM2 expression and induced cellular senescence in early postnatal primary microglia^17^ (Park et al. 2019).

The effects of excessive GCs or GR stimulation have yet to be studied in relation to human microglial development and any subsequent microglial phenotypic changes. Our study used human iPSC-derived microglia (iPS-Mg) from iPSC lines from people with distinct genetic backgrounds to construct an *in vitro* model of ELS. We report here that excessive GC at the stage of primitive haematopoiesis during microglial differentiation, resulted in increased interferon-alpha signalling, frequent formation of cGAS (a positive marker for micronuclei^18^, Miller et al. 2021), positive micronuclei indicating possible genomic instability, and senescence-like features in differentiated iPS-Mg.

## Results

The physiological receptors for GCs are glucocorticoid receptor (GR) and mineralocorticoid receptor (MR)^46^ (Timmermans et al. 2019). We determined by RNA-seq that human iPS-Mg predominately expressed glucocorticoid receptor (NR3C1), and less mineralocorticoid receptor (NR3C2) **(Figure 1B**). The NR3C1 expression level was independent of cell treatment and NR3C2 expression was not altered by treatment **(Figure 1B)**. We then determined that the predominant splice variant of the GR was GR-α, and this was present in both iPSC and iPS-Mg, whilst GR-β was not present in either the iPSC or the iPS-Mg (**Figure 1C).** Because Lieberman et al. ^47^ (2017) demonstrated that GR mRNA expression levels increased during neural differentiation, we asked whether a similar trend was observed during the differentiation of our iPS-Mg. Since GR exon 2-8 does not undergo splice variant events, and most of the GR protein isoforms originate from one species of mRNA GR^48^ (Lu and Cidlowski 2005), we used exon 7 as a readout of GR mRNA total expression. We found that there was a significant differentiation related increment in NR3C1 mRNA expression from iPSC to iPS-Mg (**Figure 1D)**

**Figure 1.**
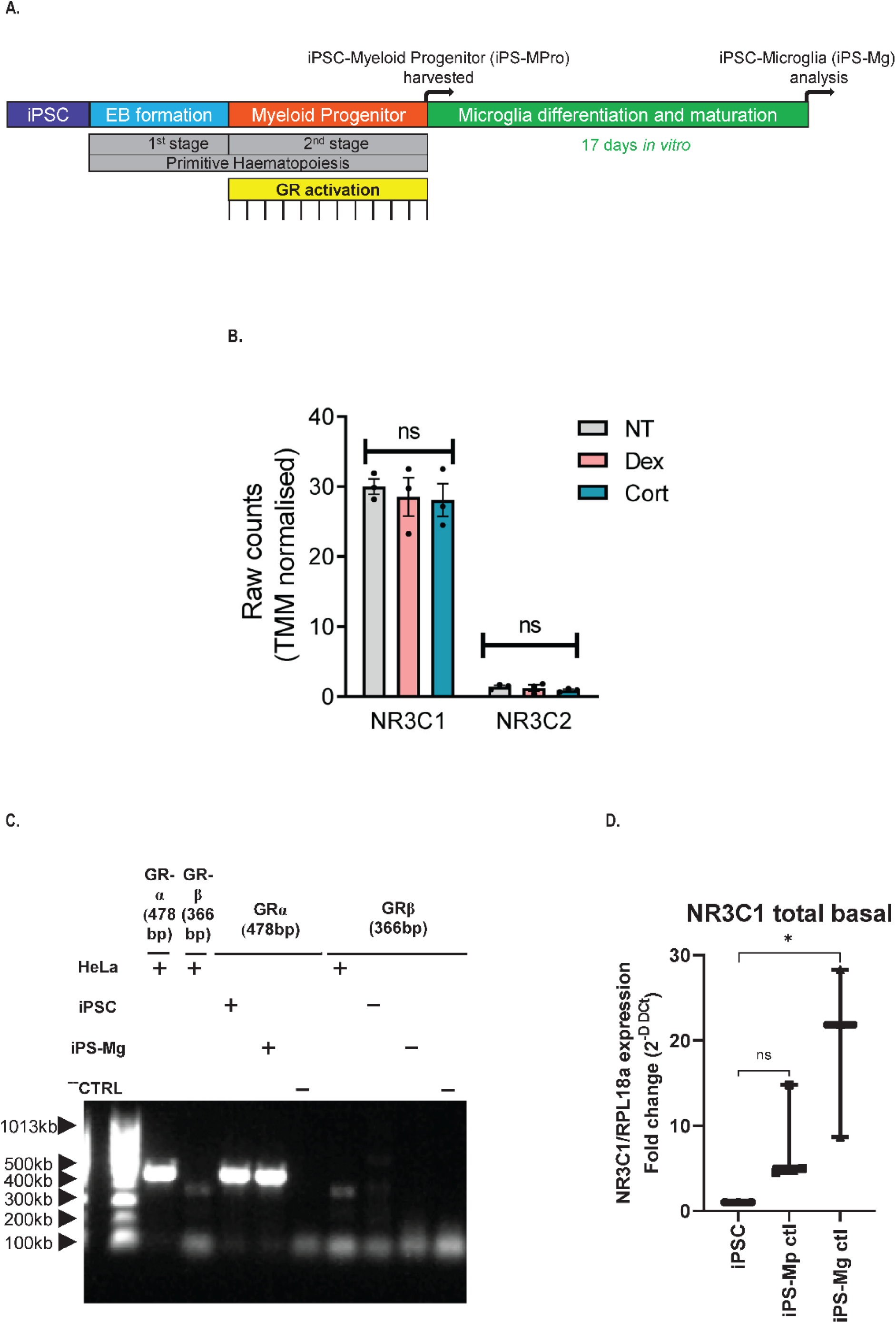
Treatment paradigm for iPSC-MPro and subsequent GR mRNA expression in iPS-Mg. **A.** Treatment paradigm for ELS via GR stimulation; the GR agonist or anatagonist (ie Dex, Cort, RU486) was added into the medium every week for approximately 12 weeks as indicated, when the medium was replenished. The iPS-myeloid progenitors (iPS-MPro) were harvested and differentiate into microglia (iPSC-Mg). **B.** Raw counts from RNA-seq analysis of GR NR3C1 and NR3C2 expressed in the iPS-Mg following no pretreatment of iPS-MPRo control cells (NT), or pretreatment with dexamethasome (Dex) or hydrocortisone (Cort). N=3. Kruskal-Wallis tests with Dunn’s multiple comparison test. Data are the mean ±SEM. **C.** iPSC and iPS-Mg expression of GR-α, and GR-β splice variant mRNA by PCR against the positive control HeLa. **D.** Expression of NR3C1 mRNA in iPSC, iPSC-MPro and iPSC-Mg. N=3. Kruskal-Wallis tests with Dunn’s multiple comparison test where *p < 0. 05. Box-and-whisker plot representing all data points and the median.

To determine the validity of using our iPS-Mg to model GR related ELS, we asked whether the microglia resembled microglia from a young age. We integrated publicly available transcriptomic data on human primary microglia from different ages across a life span of 0. 42 – 90 years of age^39,40,41,42^ (Zhang et al. 2016, Galatro et al. 2017, Gosselin et al. 2017, Olah et al. 2018) and compared this with our iPS-Mg using a hierarchical clustering method (Euclidean distance). We found that the transcriptomic profile of our human iPS-Mg clustered with young microglia **(Figure 2A).** This is in line with other papers^49, 50^ (Abud et al. 2017, Haenseler et al. 2017). These findings thus verified the use of our human iPS-Mg to model the impact of ELS at a relevant biological age.

**Figure 2.**
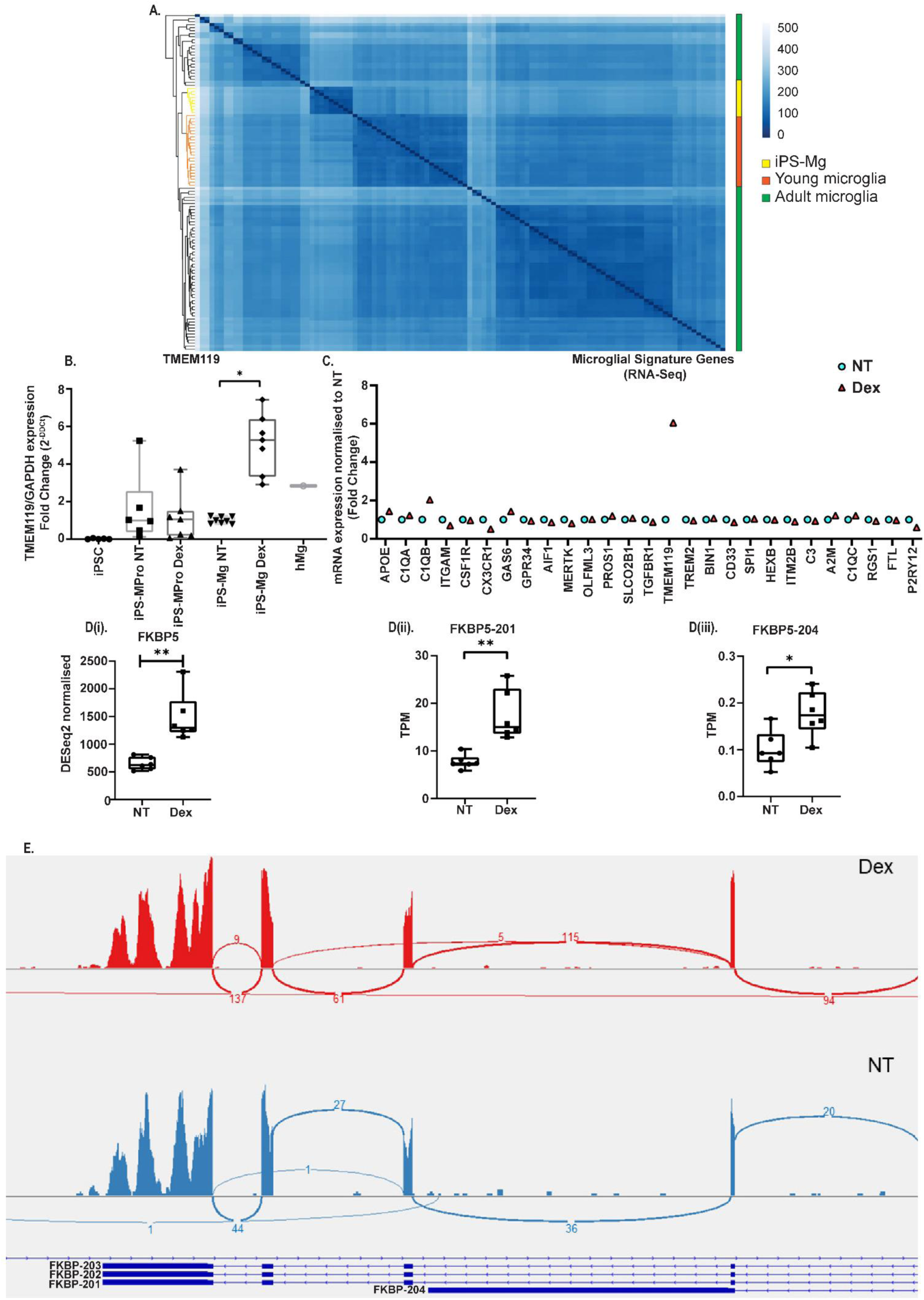
Comparative transcriptome analysis of iPS-Mg with publically - available datasets for young and adult microglia. **2A.** Heatmap of transcriptomic profile of Euclidean distance between in-house iPS-Mg and *ex vivo* primary human microglia from different ages^39,40,41,42^ (Olah et al., 2015; Zhang et al. 2016; Gosselin et al. 2017; Galatro et al. 2017). **2B**. TMEM119 mRNA expression at different cell stages from iPSC to iPSC-Mg following ELS paradigm treatment. Box-and-whisker plot representing all data points and the median, where *p < 0. 05. **2C.** RNA-seq of microglial signature genes showing increase in TMEM119 following Dex treatment. Data points are DESeq2 normalised fold change to NT from three biological replicates each condtion. **2D.** Expression of FKBP5 based on RNA-seq data analyses between NT and Dex pre-treated iPS-Mg. **i.** Total FKBP5 in NT or Dex pre-treated iPS-Mg. Values are DESeq2 normalised data from RNA-seq data. **ii.** Expression of FKBP5-201 splice variant in NT and and Dex pre-treated iPS-Mg from RNA-seq data. **iii.** Expression of FKBP5-204 splice variant in NT and and Dex pre-treated iPS-Mg from RNA-seq data. Values are from RNA-seq data and expressed as Salmon normalised TPM (transcript per million) counts. N=6. Box-and-whisker plot representing all data points and the median. Unpaired *t*-tests and *p < 0. 05, **p < 0. 01. **2E.** IGV-Sashimi plot depicting the splice variants of FKBP5 and the number of reads mapped on exons of FKBP5 between NT (blue) and Dex iPS-Mg (red).

A characteristic of a murine *in vivo* ELS model (prenatal viral infection) was the significant upregulation of TMEM119^51^ (Ozaki et al. 2020). Assessment of TMEM119 mRNA in our ELS exposed iPS-Mg mirrored this (**Figure 2B**) and was the only microglial signature gene which was upregulated (**Figure 2C).** Taken together, these data suggest that our *in vitro* model of ELS may mimic certain biological consequences associated with experimental ELS *in vivo*.

To assess whether our model of ELS can recapitulate the biological consequences of ELS *in vivo*, we conducted pathway enrichment analysis of the transcriptomic data from our ELS model and a rodent model of ELS^43^ (Delpech et al. 2016). Interestingly, the interferon alpha/beta signaling pathway and its associated pathways were found to overlap in both our ELS and an *in vivo* model of ELS^43^ (Delpech et al. 2016) (**Table 1**). Additionally, significant upregulation of FKBP5 mRNA, a co-chaperone of GRs^52^ (Binder 2009), has been observed in adult mice exposed to an adverse early life environment at an embryonic stage^53^ (Ke et al. 2018). Here the same pattern of upregulated FKBP5 mRNA was also found in our model (**Figure 2D**). Specifically, not only the total FKBP5 mRNA was upregulated **(Figure 2 Di),** but also its splice variant FKBP5-201 (**Figure 2 Dii**) and FKBP5-204 (**Figure 2D**Error! Reference source not found. **iii)**.

**Table 1.** Pathway Enrichment Analysis between our ELS model and Delpech et al. ^43^ 2016; ELS microarray gene expression data

| Pathway Enrichment | FDR |
| --- | --- |
| Interferon alpha/beta signaling | 6.11E-14 |
| Interferon alpha/beta signaling, and Interferon gamma signaling | 1.4E-13 |
| Type I interferon signaling pathway | 1.13E-12 |
| 2'-5'-oligoadenylate synthase, and type I interferon signaling pathway | 2.97E-12 |
| Mixed, incl. 4Fe-4S single cluster domain, and IFN-induced protein with tetratricopeptide repeats 3 | 0.000000193 |
| Collagen biosynthesis and modifying enzymes, and Elastic fibre formation | 0.0056 |
| **Delpech et al.<sup>43</sup> 2016: Pathway Enrichment** |  |
| Toll-like receptor signaling pathway, and Activation of IRF3/IRF7 mediated by TBK1/IKK epsilon | 9.21E-17 |
| TIR domain, and interleukin-1-mediated signaling pathway | 9.34E-10 |
| Activation of IRF3/IRF7 mediated by TBK1/IKK epsilon | 2.40E-07 |
| TNFSF members mediating non-canonical NF-kB pathway, and TNFα | 3.44E-07 |
| SMAD protein complex assembly, and c-SKI Smad4 binding domain | 5.04E-06 |
| Regulation of IFNG signaling, and interleukin-6 receptor complex | 7.12E-06 |
| Type I interferon signaling pathway, and type I interferon receptor binding | 0.00012 |
| Growth hormone receptor signaling pathway via JAK-STAT, and Leptin | 0.0128 |
| Type I interferon biosynthetic process, and Interferon beta | 0.0128 |

Sequence homology analysis was performed by using BLAST to align human FKBP5-201 transcript sequence (GRCh38) to mouse genome (GRCm39). The result yields an identity score of 85. 05% between human FKBP-201 transcript (ENST00000357266. 9), and mouse FKBP-201 transcript (ENSMUST00000079413. 11) as the top one significant alignment hit. This suggests that human and mouse FKBP5-201 have an identify score of 85. 05% and is highly conserved which facilitated the direct comparison between the expression level of FKBP5-201 in both human and mouse (**Table 2)**. The RNA-Seq of NT-iPS-Mg and Dex-iPS-Mg raw reads mapped onto FKBP-201 transcript are shown in a Sashimi plot (**Figure 2E**).

**Table 2.** Sequence homology analysis using BLAST

| Transcript ID | Transcript ID | Identities |
| --- | --- | --- |
| ENST00000357266.9 | ENSMUS 00000079413.11 | (85.05%) |

We found five of the commonly defined type I interferon (IFN)-induced genes (database from Gene Ontology) were significantly differentially expressed (DE) in the Dex pre-exposed iPS-Mg compared with NT iPS-Mg, such as the following interferon-stimulated genes (ISG) IFI6, IFI27, MX1, IFIT3, SAMD9L (adjusted ρ < 0. 1, |log2 fold-change| > 1), **(Figure 3)**. Interestingly, SAMD9L and IFI27 transcripts were significantly enriched in micronuclei containing cells^54^ (Mackenzie et al. 2017). This led us to ask if certain ISG-related signalling pathways were activated after Dex exposure and thus assessed the Dex specific upregulation of commonly defined type I IFN-induced genes on a transcriptome-wide basis. Gene set enrichment analysis (GSEA) against all genes ranked by z-scores for differential expression (Dex versus NT) calculated by DESeq2 package in R. GSEA confirmed that not only type I interferon genes are enriched in Dex-iPS-Mg (FDR = 0. 0065) (**Figure 3B)** but also innate immune response genes are enriched (FDR = 0. 0048) **(Figure 3C**).

**Figure 3.**
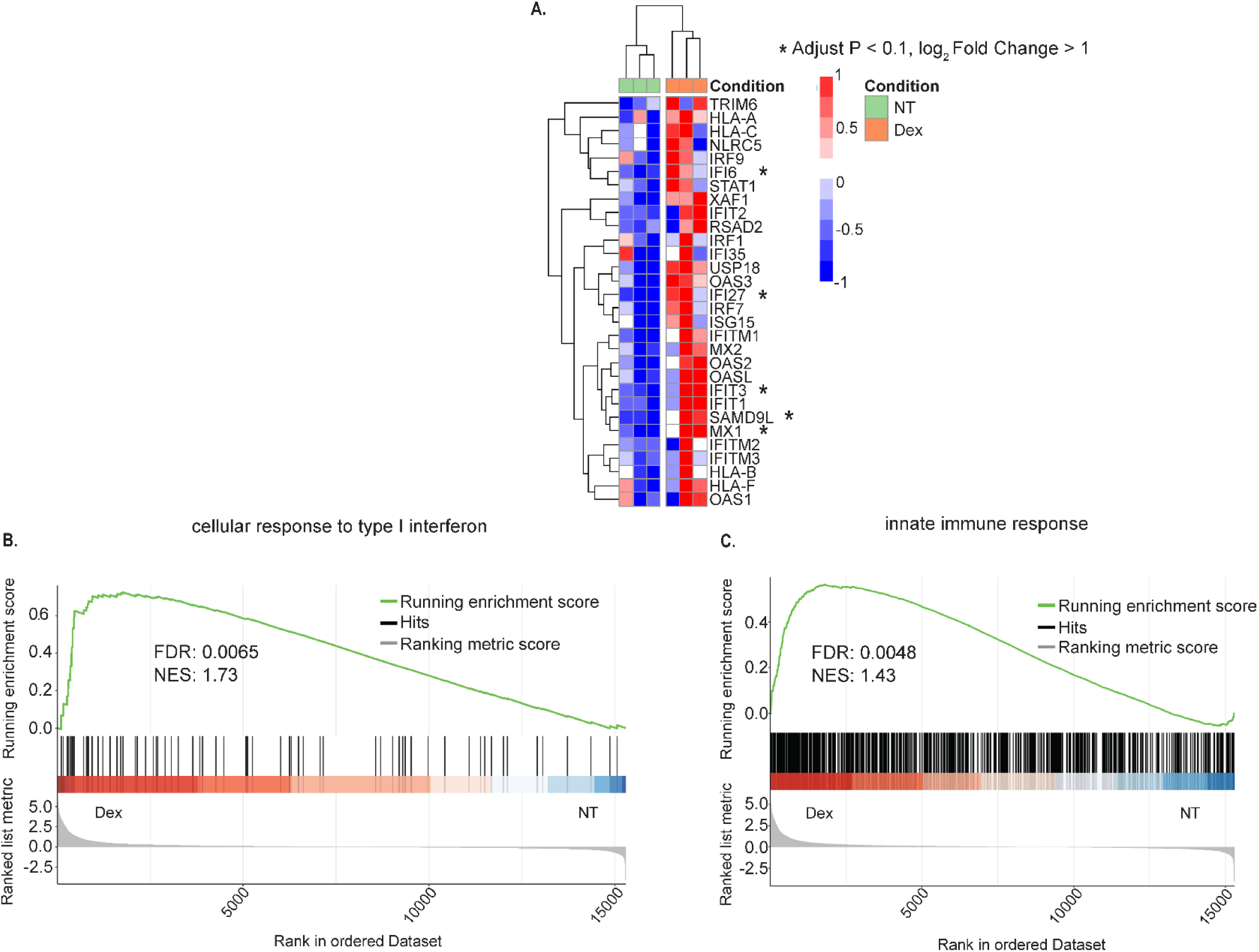
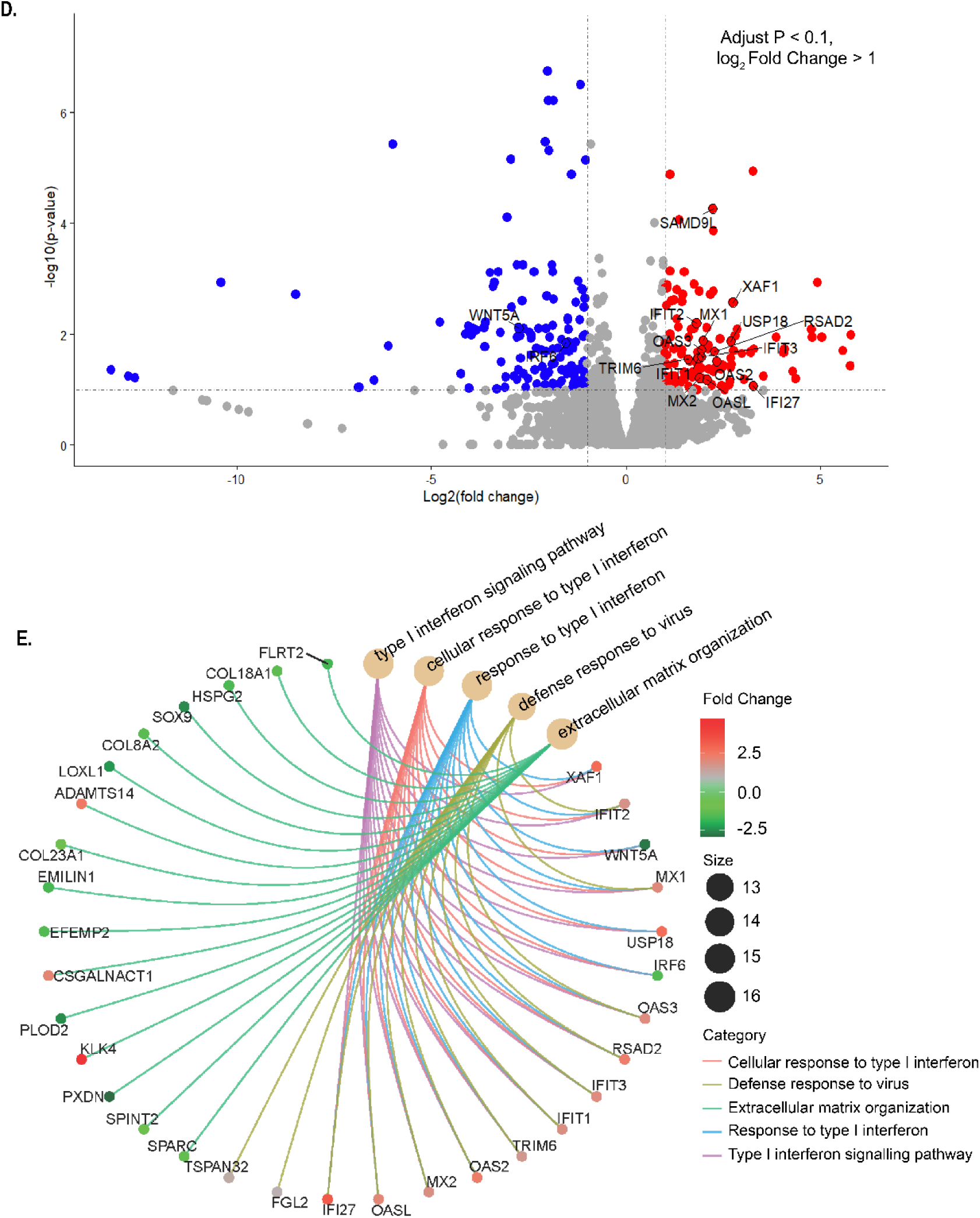
Transcriptomic analysis of Dex and Cort pre-exposed iPS-Mg. **A.** Heatmap of type I IFN-induced genes in NT and Dex pre-exposed iPS-Mg. DESeq2 normalised data are plotted. * adjusted p < 0. 1, |log2 fold-change| > 1. **B., C.** Transcriptome-wide analysis (GSEA) of commonly defined type I IFN-induced genes (FDR =0. 0065) (**B**) and innate immune response genes (FDR = 0. 0048) **(C)** are significantly enriched in transcriptomes of Dex pre-exposed iPS-Mg. **D**. Volcano plot illustrating differential expression of genes in Cort pre-exposed iPS-Mg and NT iPS-Mg. Genes with adjusted p < 0. 1, and |log2 fold-change| > 1 are plotted in red and blue. Red indicates upregulation, blue indicates downregulation. Type I IFN-induced genes were circled. **E.** Gene ontology analysis of the significantly differentially expressed genes, and significantly enriched signalling pathways in Cort pre-exposed iPS-Mg compared with NT iPS-Mg.

To confirm these findings, an endogenous agonist of GR, hydrocortisone (Cort) was used in the same way as Dex treatment. Analysis of Cort-iPS-Mg revealed 125 significantly upregulated and 129 downregulated genes compared with NT iPS-Mg (adjusted p < 0. 1, |log2 fold-change| > 1, (**Figure 3D**). Consistent with Dex-iPS-Mg, Gene Ontology analysis on the significantly DE genes affirmed activation of type I interferon signalling in Cort-iPS-Mg (**Figure 3E**).

To further validate the activation of type I interferon signalling when iPS-Mg were pre-exposed to GR, we tested our hypothesis on a different genetic background. Patient-derived iPS-Mg expressing the Alzheimer’s Disease (AD) risk variant R47H^het^ in triggering receptor expressed on myeloid cells 2 (TREM2 R47H^het^) were treated with Dex in the same manner as our control lines used above (SFC840, BIONi010-C, and KOLF2_C1)^19, 22^ (Cosker et al. 2021; Piers et al. 2020). To gain an understanding of obvious and subtle changes, we performed weighted gene co-expression network analysis (WGCNA). We integrated RNA-Seq of control lines and TREM2 variant lines +/- Dex, Cort into the WGCNA analysis (15 samples in total) and detected a total of 32 modules **(Supplementary Figure 1i)**.

For module-trait analysis, we categorised Dex and Cort and NT as the GR activation trait. Firstly, this is because our previous analysis indicated that Dex and Cort treatment led to activation of similar pathways **(Figure 3**); secondly, *in vitro* studies of the pharmacological characteristics of Dex and Cort in cells expressing endogenous GR show that the concentrations of Cort or Dex used in the present study are at the receptor EC_50_ ^55^(Longui et al. 2005), which facilitates the combination of Cort and Dex, a GR activation trait for WGCNA.

Out of 32 detected modules, the module eigengene of 7 modules, namely royalblue, darkturquoise, green, red, darkgrey, yellow and purple were significantly correlated with the GR activation trait **(Figure 4A, marked with**\*). Also, these 7 GR modules are not affected by the genetic variation between the control lines used here (which express the common TREM2 variant) and the R47H^het^ TREM2 variant lines.

**Figure 4.**
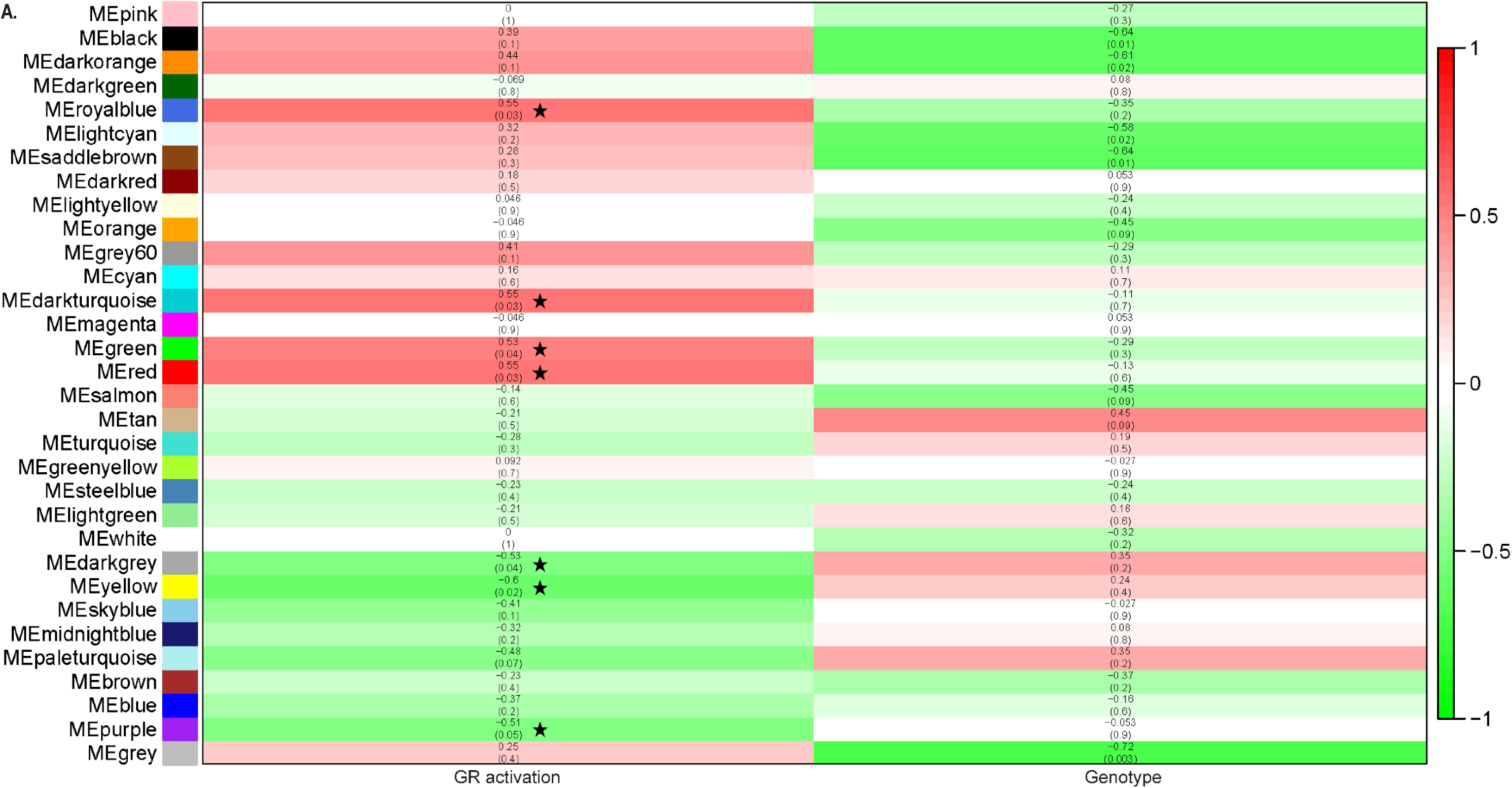
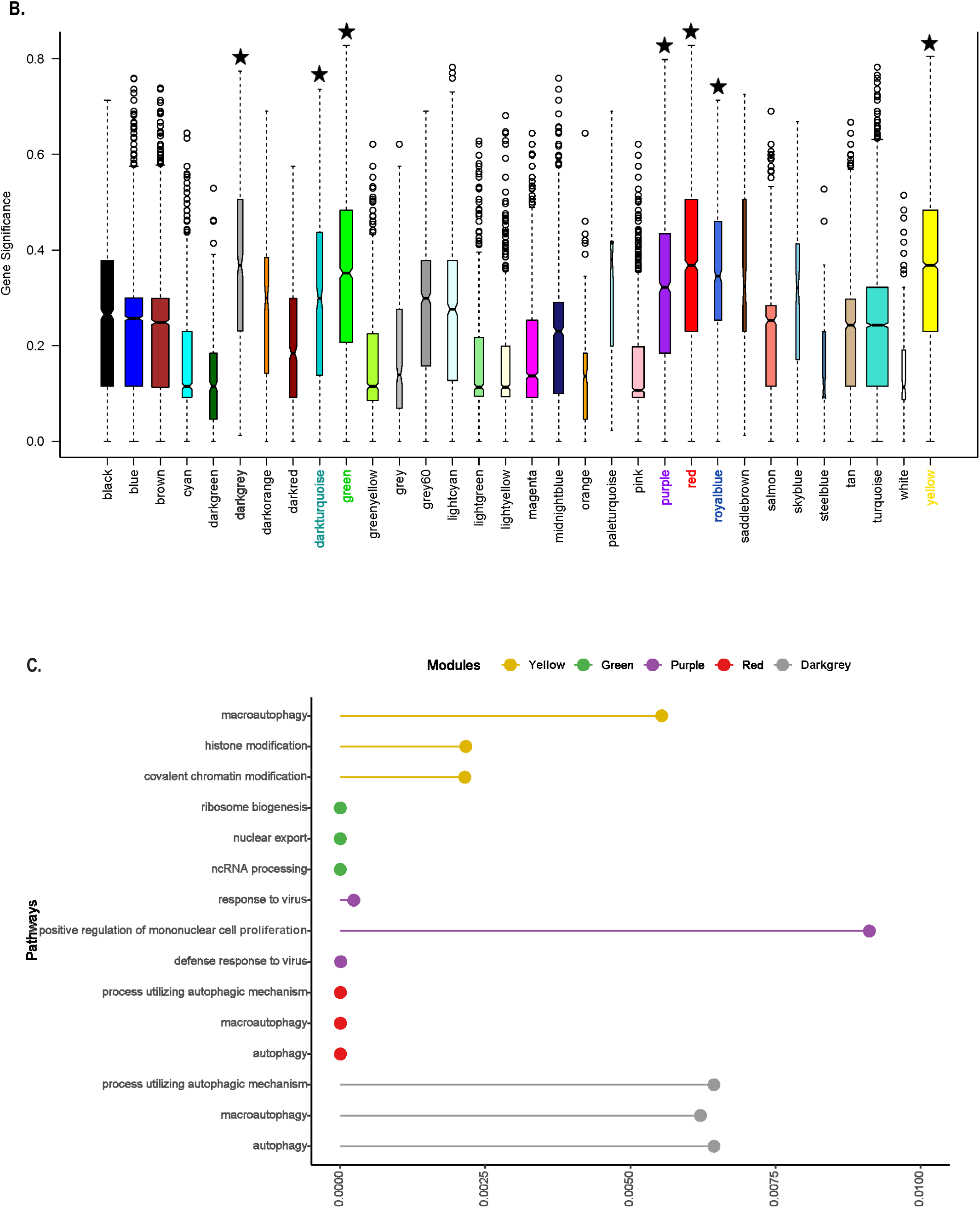
WGCNA analysis of Dex and Cort pre-exposed iPS-Mg with Cv and TREM2 variants. **A.** Diagram of module-trait relationship reports Kendall’s correlation coefficients (shown at the top of each row), and its corresponding p values (displayed at the bottom of each row within parentheses) between the eigengene value of each module, two traits GR activation and genotype are displayed at the bottom of the diagram as column. Modules specifically related to GR activation and were not correlated with genotype are marked with star. The direction of the correlation was color-coordinated with positive correlations indicated in red and negative correlation in blue. **B.** Distribution of average gene significance in the modules associated with GR activation trait. The 7 GR activation related modules previously identified (marked with star) showed higher overall gene significance value than other modules and were selected for further analysis. **C.** GO enrichment analyses of genes in yellow, green, purple, red and darkgray modules, top 3 significantly enriched pathways in each pathway were plotted. Bars representing GO terms show Benjamini and Hochberg FDR adjusted p values (q. value).

By using gene significance (GS) and module membership (MM) analysis, we ensured that the GR modules are specific to GR activation traits (**Figure 4B and Supplementary Figure 1ii & 1iii).** To ensure the GR modules are high quality modules, we implemented module preservation analysis on one single topological overlap matrix (TOM) network both as test and reference. Since only one TOM was constructed, module quality was assessed in the present study. We calculated multiple module quality statistics that measure how well-defined modules are in repeated random splits of the reference dataset. All modules demonstrated strong evidence for high quality (Zsummary > 10), confirming that GR modules were well-defined and non-random **(See Supplementary Figure 2A).**

To gain a deep understanding of what GR pre-treatment does to iPS-Mg, we further analysed the GR activation-related modules. Since genes that are highly co-expressed often share similar functions, biological processes and pathways that are enriched in a co-expression module can be used to infer functional information^56, 57^ (Singer et al. 2004, de la Fuente 2010). We performed Gene Ontology enrichment analysis on the 7 identified GR modules to detect significantly overrepresented gene ontology categories in each WGCNA module. The top three significant pathways in each module were plotted (**Figure 4C**). The purple module is enriched in pathways associated with defence responses to viruses (q. value = 7. 59E-06), which confirmed the type I interferon signalling pathway identified in our DE gene analysis. The darkgray (q. value = 0. 00553918), red (q. value = 2. 45E-06) and yellow modules (q. value = 0. 006206759) are significantly enriched with genes associated with macro-autophagy and the autophagy pathway. The green module is enriched with pathways associated with ribosome biogenesis (q. value = 2. 00E-25), nuclear export (q. value = 4. 41E-19) and non-coding RNA processing (5. 62E-22 q. value). Darkturquoise and royalblue yielded no significant pathways within the modules.

To follow up on the finding of enhanced type I interferon and associated viral and viral defence signalling in Dex-iPS-Mg, we reasoned that one possible cause of this signal might be genomic self-DNA in the cytoplasm^58^ (Keating et al. 2011). In this ectopic location, the self-DNA would be perceived as an invading virus and trigger an anti-viral defence and transcription of type I interferon^59^ (Honda et al. 2006).

We investigated the possible presence of cytosolic self-DNA. Following crude cytosolic/nuclear fractionation, we found that Dex-iPS-Mg displayed significantly more cytosolic self-DNA compared with NT-iPS-Mg or RU/De-iPS-Mg (**Figure 5A, Supplementary Figure 2B**) and that this was prevented by RU treatment (**Figure 5Aii**).

**Figure 5.**
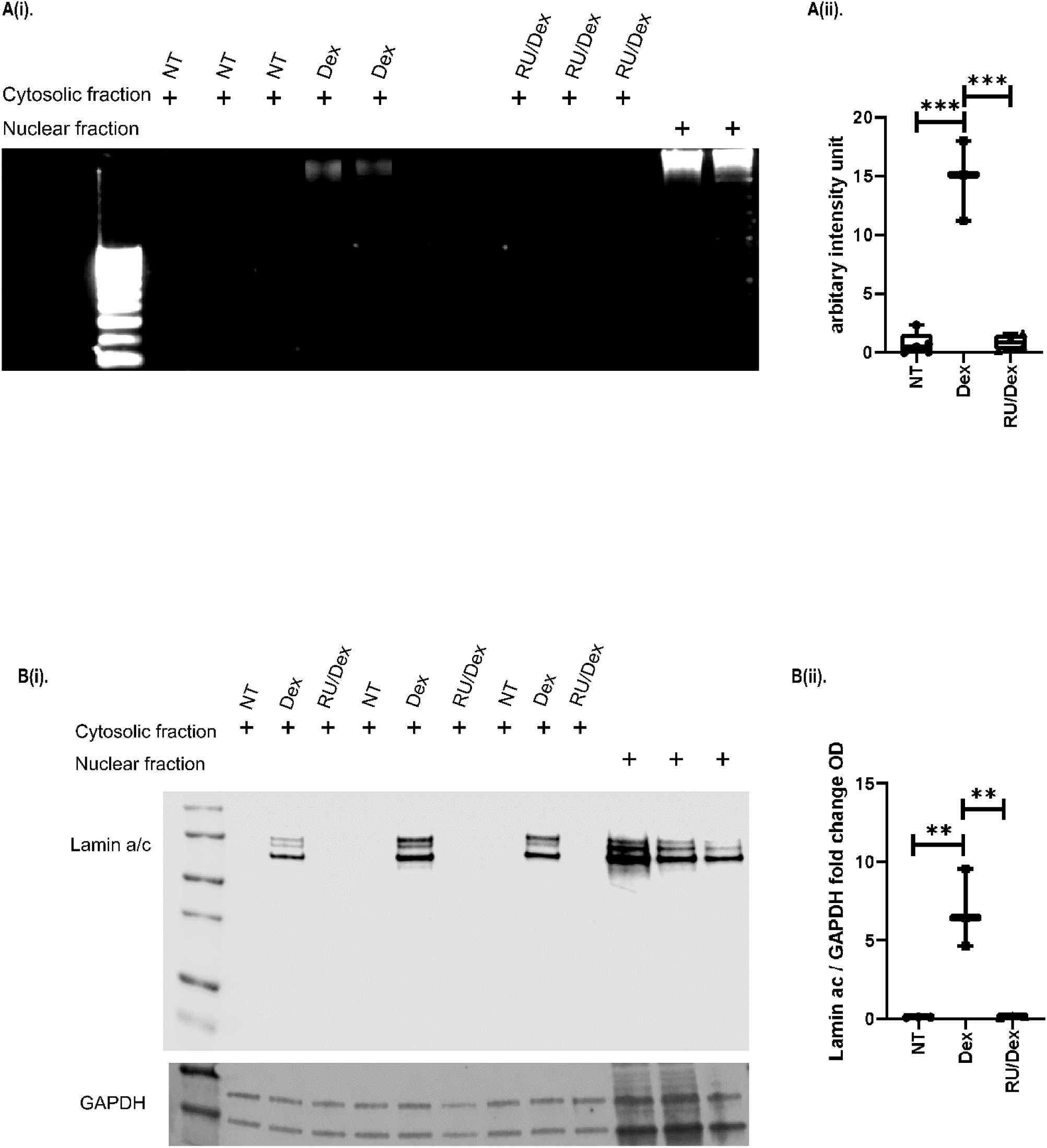
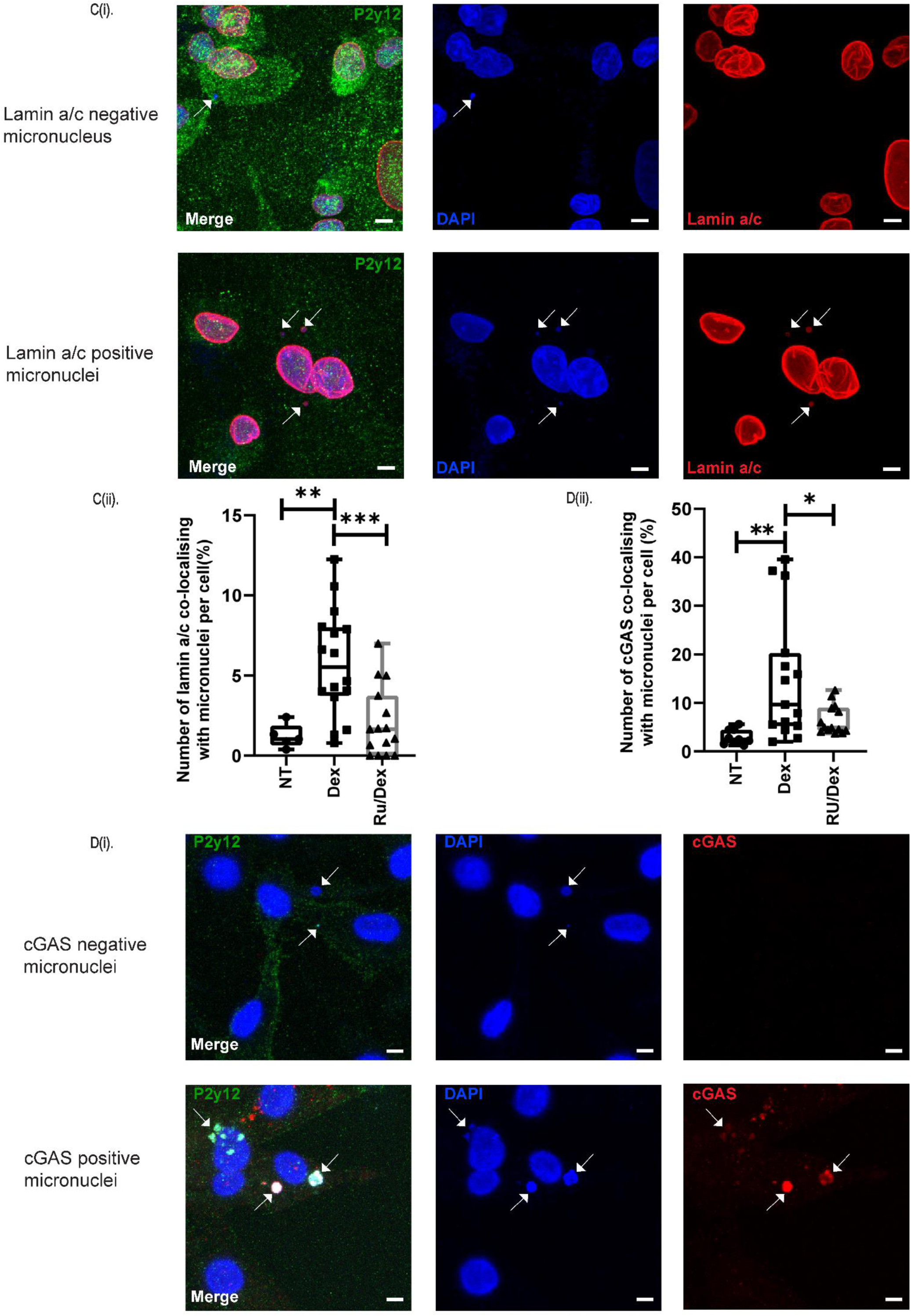
Formation of micronuclei and cGAS activation in Dex pre-exposed iPS-Mg. **Ai.** Agarose gel analysis of genomic DNA in cytosolic or nuclear extracts from NT iPS-Mg, Dex pre-treated iPS-Mg and RU/Dex pre-treated iPS-Mg. **A ii.** Quantification of the genomic DNA present in the cytosolic and nuclear fraction from the gel shown in **Ai**. N = 3 biological replicates (additional replicates are shown in Supplementary Fig 1C). One-way ANOVA with Dunn’s multiple comparison test. Box-and-whisker plot representing all data points and the median. **Bi.** Western blot of Lamin a/c in cytosolic and nuclear fractions. Nuclear fractions are shown as positive controls for Lamin a/c. GAPDH served as a loading control in the cytosolic extract. **B ii.** Quantification of Western blot shown in Bi. N=3 One-way ANOVA with Dunn’s multiple comparison test. Box-and-whisker plot representing all data points and the median. **Ci.** Immunostaining with antibody against Lamin a/c (red), P2y12 (green) with DAPI nuclear counterstain (blue) in NT iPS-Mg, Dex pre-treated iPS-Mg and RU/Dex pre-treated iPS-Mg. Scale bar: 5µm. **Cii.** Number of micronuclei/cell in NT iPS-Mg, Dex pre-treated iPS-Mg and RU/Dex pre-treated iPS-Mg. N=3 biological replicates. One-way ANOVA with Dunn’s multiple comparison test. Box-and-whisker plot representing all data points and the median. Each data point represents one whole field of 10X10 tile scan per condition. Arrow points to lamin a/c positive and negative micronuclei. **Di.** Immunostaining with antibody against cGAS (red), P2y12 (green) with DAPI nuclear counterstain (blue) in NT iPS-Mg, Dex pre-treated iPS-Mg and RU/Dex pre-treated iPS-Mg. Scale bar: 5µm. **Dii** Number of cGAS colocalising with micronuclei per cell in NT iPS-Mg, Dex pre-treated iPS-Mg and RU/Dex pre-treated iPS-Mg. N= 3 biological replicates, one-way ANOVA with Dunn’s multiple comparison test to compare nontreated group with the Dex and RU/Dex pre-treated group, *p < 0. 05; **p < 0. 01; ***p < 0. 005. Box-and-whisker plot representing all data points and the median. Each data point represents one whole field of 10X10 tile scan. Arrow points to cGAS positive and negative micronuclei.

To ensure that the cytosolic extract was not contaminated by spill-over from the nuclear fraction, the cytosolic extract was probed for lamin a/c, a nuclear protein^60^ (Dubik and Mai 2020). We found in NT-iPS-Mg, there was no lamin a/c, which confirmed that the cytosolic extract was not contaminated by the nuclear fraction. Conversely, there was significantly more lamin a/c in Dex-iPS-Mg, which was attenuated by pretreatment with GR antagonist RU486 (**Figure 5Bi & ii**).

This finding led us to speculate that the cytosolic self-DNA with lamin a/c could be micronuclei^61^ (Kneissig et al. 2019). Micronuclei are lamin a/c positive small DNA-containing nuclear structures that are spatially isolated from the main nucleus^62^ (Kwon et al. 2020). Confocal imaging analysis revealed that Dex-iPS-Mg displayed significantly more micronuclei per cell compared with NT-iPS-Mg or RU/Dex-iPS-Mg (**Figure 5C**). To ensure that we do not mistake apoptotic bodies as micronuclei, we ensured that pre-treated of Dex or RU486 alone do not induce significant cell death (**Supplementary Figure3 A & B**).

Micronuclei are susceptible to nuclear envelope collapse^63^ (Hatch et al. 2013) due to an unstable micronuclei envelope. The DNA in the micronuclei can serve as a source of immunostimulatory cytosolic DNA. Following this, cyclic GMP-AMP synthase (cGAS) binds to self-exposed DNA in the micronuclei, 2’-5’-cGAMP is synthesized, leading to the activation of stimulator of interferon genes (STING) and transcription of type I interferon^64^ (Zierhut and Funabiki 2020). Since cGAS is upstream of this pathway, and cGAS localises to micronuclei upon nuclear envelope rupture^54^ (Mackenzie et al. 2017), we investigated the number of cGAS colocalised micronuclei per cell in NT-, Dex- and RU/Dex-iPS-Mg. We detected a significantly higher number of cGAS positive micronuclei in Dex-iPS-Mg compared with NT-iPS-Mg (**Figure 5D**), or RU/Dex-iPS-Mg.

One observation of GR activation, was a significant reduction in the cell number of Dex-iPS-MPro compared with NT-iPS-Mg or RU/Dex-iPS-Mg (**Figure 6A**). This suggested that certain proliferation related processes or cellular components might be affected by Dex exposure. Thus, we revisited the GSEA analysis on the whole transcriptomics of NT and Dex-iPS-Mg, specifically focusing on the top 10 highest score of gene ratio among all other examined enriched pathways. Gene ratio is the percentage of total DEGs in the given GO term. We discovered that the condensin complex (cellular component, CC), mitotic spindle elongation and spindle midzone assembly (biological processes, BP) are all suppressed (**Figure 6B, marked with** *). Taken together, suppressed genes of condensin complex and mitotic spindle elongation, and significantly reduced cell number in Dex-iPS-Mg suggests defective proliferation, and possibly cellular senescence.

**Figure 6.**
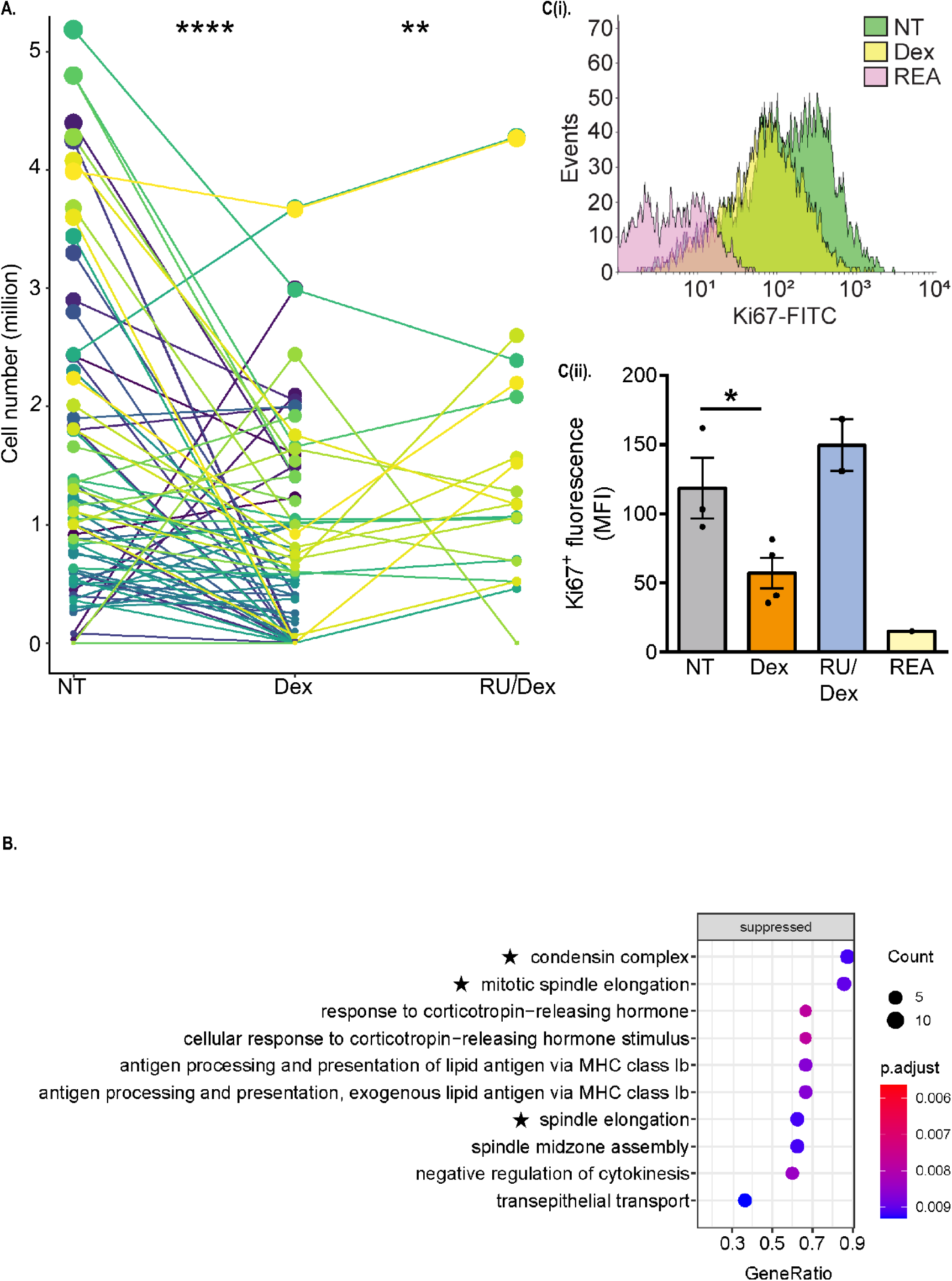
Dex-treated iPS-MPro and pre-treated iPS-Mg exhibit reduced cell number and proliferation. **A.** Before-after graph of the iPS-MPro cell number generated during primitive haematopoiesis either non-treated (NT) or treated with Dex or Ru/Dex. Lines connected with dots across treatments mean one collection event. Kruskal-Wallis test with Dunn’s multiple comparison test of NT versus Dex (****p < 0. 0001) or Dex versus RU/Dex (**p < 0. 01). **B.** Dot plots of GSEA results illustrating top 10 GO biological processes and cellular components associated with consequences of Dex-pretreated iPS-Mg based on gene ratio. ‘Gene ratio’ is the percentage of total DEGs in the given GO term (only input genes with at least one GO term annotation were included in the calculation). **Ci** Representative FACS histogram of cells stained with anti-Ki67 antibody in NT, Dex and RU/Dex pre-treated iPS-Mg. **Cii.** Quantification of Ki-67 protein expression as mean fluorescent intensity (MFI) shown in C. Data are the mean of n= at least 3, analysed with Two-tailed unpaired t test. *p < 0. 05. Data are the mean with ±SEM.

The link between cGAS positive micronucleated cells and senescence is well established^65^ (Glück and Ablasser 2019). Since Dex-iPS-Mg expressed an increase in the number of cGAS positive micronuclei, the microglia might also exhibit cellular senescence. As one characteristic of senescent cells is the loss of proliferative capacity^66^ (Galvis et al. 2019), we analysed the expression of the proliferation marker, Ki67 in NT-,Dex- and RU/Dex-iPS-Mg. Dex-iPS-Mg displayed a significantly reduced Ki67 expression compared with NT, and relatively lower expression of Ki67 compared with RU/Dex (**Figure 6C**).

We also examined changes to three “gold standard" senescence markers, namely, senescence-associated beta galactosidase (SA-β-gal), senescence-associated Secretory Phenotype (SASP), and relative telomere length (reviewed in Galvis et al. ^66^ 2019). Dex treated iPS-Mg showed significantly greater intensity of SA-β-gal staining compared with NT or RU/Dex iPS-Mg (**Figure 7A)**. In addition, Dex-iPS-Mg displayed a trend towards shorter telomere lengths compared with NT-iPS-Mg (**Figure 7B).** Furthermore, of 105 cytokines and chemokines investigated as secreted in conditioned medium, the most significantly upregulated were 11 SASP proteins previously shown to be upregulated in senescent cells^67^ (Lunyak et al 2017) and here showed increased expression in Dex compared with NT or RU/Dex-iPS-Mg **(Figure 7C).** Taken together, we demonstrated here that in Dex-iPS-Mg, there is an overarching presence of senescent characteristics.

**Figure 7.**
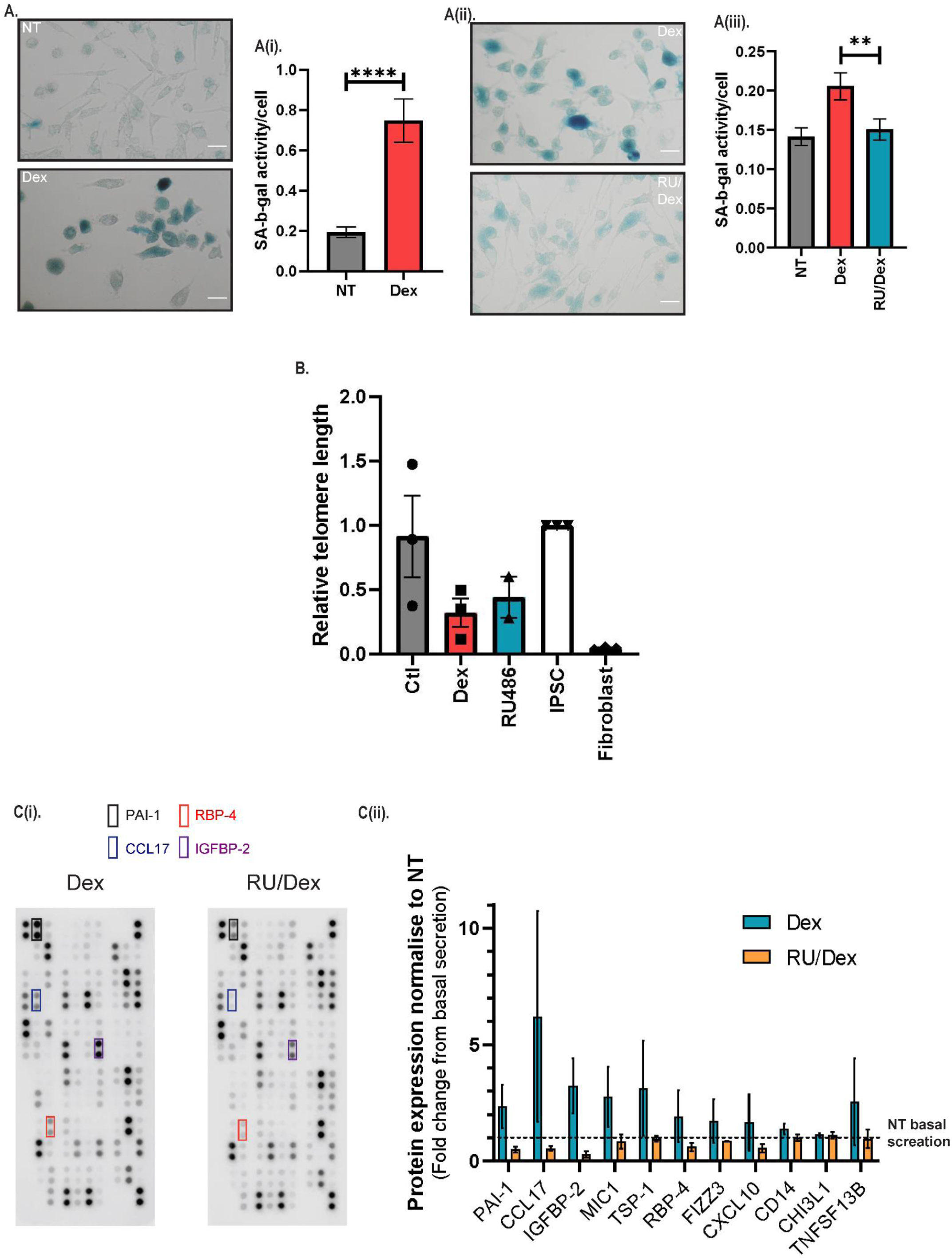
Dex-pre-treated iPS-Mg exhibit characteristics of cellular senescence. **A.** Representative images of senescence-associated beta galactosidase (SA-β-gal) staining in NT and Dex pre-treated iPS-Mg. **Ai.** SA-β-gal activity quantified by staining intensity in NT, Dex pre-treated iPS-Mg normalised to cell number. N=3, Mann-Whitney test. **** p < 0. 0001. Data are the mean ±SEM. **Ai.** Representative images of senescence-associated beta galactosidase (SA-β-gal) staining in Dex and RU/Dex pre-treated iPS-Mg. **Aii.** SA-β-gal activity quantified by staining intensity in Dex or RU/Dex pre-treated iPS-Mg normalised to cell number. N=4. Kruskal-Wallis test with Dunn’s multiple comparison test. **p < 0. 01. Data are the mean ±SEM. **B.** Relative telomere length determined by real-time qPCR in NT, Dex and RU/Dex pre-treated iPS-Mg. N=3. **Ci.** Representative human XL cytokine proteome array dot blots from Dex and RU/Dex pre-treated. N=2 membranes with 2-3 separate pooled replicates. **Cii.** Quantification of the expression of 11 Senescence-Associated Secretory Phenotype (SASP) related proteins. Expression level of SASP related proteins are normalised to NT iPS-Mg as basal secretion. Dex treated iPS-Mg has overarching higher expression of SASP related proteins. N=2. Data are the mean ±SEM. Statistical significance was addressed using one way ANOVA with Dunnett’s multiple comparison test, or Kruskal-Wallis test with Dunn’s multiple comparison test to compare nontreated group with the Dex and RU/Dex treated group, *P < . 05; **P < . 01; ***P < . 005

## Discussion

Here, we demonstrate for the first time that human iPSC-derived microglia express GR, and a developmental- and differentiation-related increase in GR mRNA expression. Furthermore, prolonged activation of GR during human iPS-Mg differentiation triggered enhanced signalling of type I interferon, formation of cGAS positive micronuclei together with cellular senescence in the fully differentiated and maturated iPS-Mg. As we only modulated GR activation during primitive haematopoiesis before differentiation and maturation of iPS-Mg, what we observed and reported here is the priming effect and after-effect of aberrant GR activation during microglial development.

Our finding that the transcriptomic profile of our human iPS-Mg mostly clustered with young microglia is in line with previous papers^49, 50^ (Abud et al. 2017, Haenseler et al. 2017). With regards to the GR expression, we show that human iPSC-derived microglia express GR NR3C1, and furthermore that this is the splice variant GR-α. This corresponds to the active form of the receptor, ie one that can be activated by cortisol^68^ (Hagendorf et al. 2005). GR-α levels and activity are modulated by alternative splicing of the common pre-mRNA into mRNAs for the GR-β and GR-P isoforms^68^ (Hagendorf et al. 2005). We did not detect the expression of GR-β in our cells, either at the iPSC stage or following differentiation to microglia. Our finding that GR activation (by Dex) led to a reduction in cell number (of iPS-MPro) has been reported for other cell types^69^. ^70^ (Samuelsson et al. 1999, Piette et al. 2009), where it reflected an inhibition of proliferation^70^ (Piette et al. 2009). In line with this, we report that proliferation (as measured by Ki67 expression) was reduced in the iPS-Mg following their early exposure to Dex at the iPS-MPro stage (suggesting a maintained phenotype effect). We also found that from the development of the iPSC to iPS-Mg, the expression of NR3C1 increased. Previously Lieberman et al. ^47^ (2017) demonstrated that GR mRNA expression levels increased during the differentiation of iPSC-derived neural cells; GR mRNA has also been shown to be increased in human iPS derived cerebral organoids^71^ (Cruceanu et al. 2020). We also report that GR activation led to an increase in FKBP5, FKBP5-201 and FKBP5-204. FKBP5 is the intrinsic regulator of GR activity and has been shown to promote inflammation by activating the master immune regulator NF-κB, and increasing the release of proinflammatory cytokines, such as IL-8^72^ (Zannas et al. 2019). This is in accord with our findings, where we found that GR activation led to enhanced type I interferon signalling as detailed in our RNA-seq data, a driver of antiviral response pathways in microglia.

The differentiation/maturation of iPS-Mg follows the cellular ontogeny of microglia (primitive haematopoiesis, microglial differentiation and maturation with signals from other CNS cells. It is therefore possible to introduce GC mediated stress before microglia are differentiated and matured as a model of ELS. With staged differentiating iPS-Mg, we activated GR with 50 nM dexamethasone (Dex) treatment in the medium during myeloid progenitor generation. We choose to activate GR at the stage of primitive haematopoiesis, because it is the first critical stage for microglial development^15^ (Ginhoux et al. 2010). Following that, myeloid progenitors were harvested for microglial differentiation and maturation without any treatment. Therefore, the present study investigated the consequences after chronically activating GR at primitive haematopoiesis; specifically, the priming effect of aberrant GR activation at primitive haematopoiesis on differentiated/matured iPS-Mg, from a developmental perturbation angle. Accordingly, the present study utilized a Dex concentration of 50 nM, which is sub-therapeutic and within a physiological relevant range^73^ (Jameson et al. 2006); human preterm infants typically receive 0. 1-0. 15 mg/kg Dex, which is about 0. 6 µM when given directly to a preterm infant, or 0. 06 µM in the foetus when Dex is given to the mother^74, 75^ (Osathanondh et al. 1977, Charles et al. 1993). The use of the endogenous GR activator, hydrocortisone (Cort), at a comparative concentration to Dex treatment^55^ (Longui et al. 2005), confirmed the findings of the synthetic agonist.

Previous models investigating the biological effect of ELS captured only the immediate effect on embryonic or early postnatal microglia following short ELS exposures. Thus, prenatal elevated GC was shown to increase overall microglial density and promote an immunoreactive embryonic microglial phenotype following short 2 days ELS exposure^16^ (Bittle and Stevens 2018) which differs from our findings. A 72-hr aberrant GR activation in early postnatal rodent primary microglia was found to downregulate CX3CR1 and TREM2 expression and induce a senescence phenotype^17^ (Park et al. 2019). Our present study provides insight into the priming effect of early chronically activated GR on subsequent microglia development.

Other models of ELS, such as maternal immune activation (MIA) in mice, found disrupted placental GC metabolism^76^ (Núñez Estevez et al. 2020). The compromised placental GC metabolism could result in dysregulated levels of GC bypassing the placenta and entering the foetus, and leading to aberrant activation of foetal GR. It is increasingly evident that MIA has a significant impact on certain aspects of microglial biology. For example, MIA-exposed young microglia become more pro-inflammatory and exhibited a different morphology compared with young microglia without MIA exposure^77, 78^ (Diz-Chaves et al. 2012, Diz-Chaves et al. 2013); and MIA induced by proinflammatory cytokines led to alterations in foetal microglial tip motility patterns and increased mRNA expression of TMEM119^51^ (Ozaki et al. 2020) mirroring our finding of increased TMEM119 mRNA expression. Although the effect of MIA on microglia might be the primary reason for the altered microglial biology, it is likely that dysregulated maternal GC and subsequent aberrant foetal GR activity could co-occur within *in vivo* models of MIA and would need further investigation.

One possible explanation of our treatment paradigm of GCs and the resulting phenotype we report, may be that ELS mediated excessive GC results in the persistent occupation of the ligand-binding domain of GRs. This results in less unoccupied GRs and in a subsequent loss in the ability to accurately regulate chromosome segregation. This is because GRs are critical for the regulation of accurate chromosome segregation. Specifically, Matthews et al. ^79^ (2015) pointed out that the ligand-binding domain of unliganded (free) GRs plays a direct role in regulating mitotic spindle function. This is in line with our finding of a higher number of micronuclei in Dex pre-treated iPS-Mg, and a lower number of micronuclei when GR activation was blocked. However, our finding contrasts to the classical anti-inflammatory and immunosuppressive properties of acute GC treatment, where GCs have an inhibitory effect on the type I interferon signalling pathway^80^ (Flammer et al. 2010). Taken together, our current findings add to the novel property of GCs in the context of microglial development.

To date, research findings of ELS via GR on microglia have primarily come from rodent studies. Moreover, it has proven difficult to translate findings from rodent microglia to human microglia. This is because there are different immune responses between humans and animal models including the fact that experimental rodents are reared in contained environments, and exhibit life-span differences^81^ (Smith and Dragunow 2014). Given these caveats, the current model can provide an alternative model for manipulation at an early developmental stage which has an impact on subsequent microglial phenotypes.

Here we confirmed that one possible cause of enhanced type I interferon and viral defence signalling in Dex-treated iPS-Mg was a more frequent formation of micronuclei with activated cGAS. Thus, the cytosolic presence of self-DNA with lamin a/c was specifically dependent on aberrant activation of GR during primitive haematopoiesis in iPS-Mg. Increased formation of cGAS positive micronuclei is found in cancer cells^82^ (Gisselsson et al. 2001), and indicates genomic instability^54^ (Mackenzie et al. 2017). To our knowledge, this is first time that prolonged activation of GR during primitive haematopoiesis results in genomic instability and cellular senescence in the resultant microglia. Our findings have implications for the aetiology of neurodevelopmental disorders, such as ASD and schizophrenia. For example, recent evidence points to genomic instability and copy number variation in ASD and schizophrenia^83^ (Kushima et al. 2018). This is in line with our current findings. In particular, the phenomenon of genomic instability shown here in human iPS-Mg is relevant to these conditions as microglia are a CNS cell type heavily engaged in neurodevelopment.

In summary, we have established that an *in vitro* model of ELS in iPS-Mg is able to induce certain biological consequences of ELS found *in vivo*. We found prolonged activation of GR during microglial development leads to genomic instability via frequent formation of cGAS positive micronucleus and complemented with cellular senescence. These data have ramifications for developmental conditions following maternal stress.

## Acknowledgements

The running costs for the project were provided by J. M. P and J. H. . Additional funding for the RNA-seq analysis was provided to J. W., T. M. P. and J. M. P. with a pump-priming grant from the Alzheimer’s Research UK. This research used the UCL genomics facilities and UCL Cancer Institute sequencing facilities. C. A. was supported by a fellowship from the Alzheimer’s Society UK (AS-JF-18-008) in the lab of S. W. who was supported by an Alzheimer’s Research UK Senior Research Fellowship (ARUK-SRF2016B-2) and the National Institute for Health Research University College London Hospitals Biomedical Research Centre. We acknowledge access to the database of Genotypes and Phenotypes (dbGaP) for datasets on young microglia (Gosselin et al. 2017), mid-age microglia (Zhang et al. 2016), and SYNAPSE (Olah et al. 2018) for datasets on old microglia, and all age microglia (Galatro et al. 2017).

## Contributions

J. W. and T. M. P. devised the project with input from J. M. P. J. W. carried out all the experiments. The pathway enrichment analysis was done by T. M. P. C. H. assisted with the qPCR, J. W. and J. M. P. wrote the paper. All authors reviewed the paper.

## Figure Legends

**Supplementary figure 1i.**
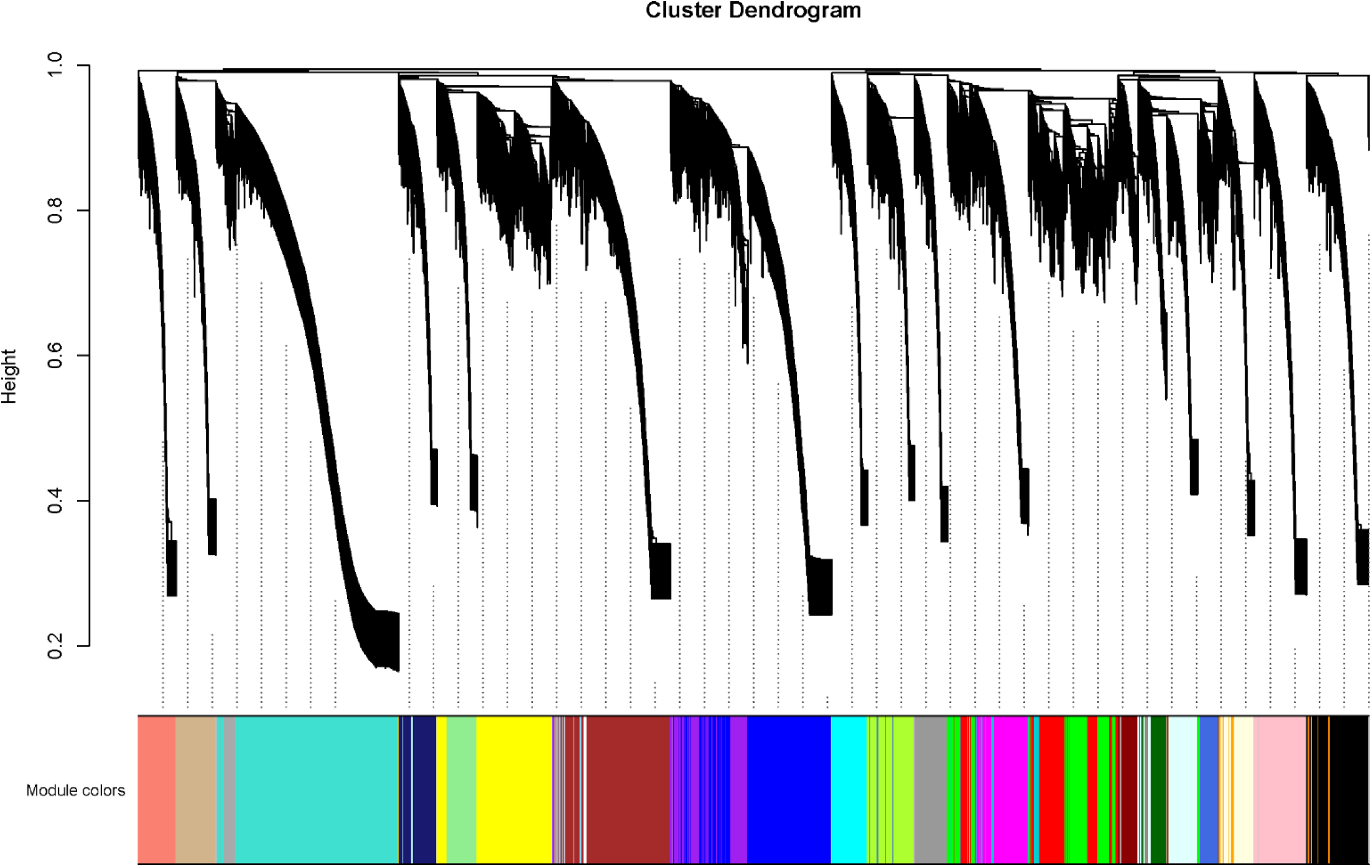
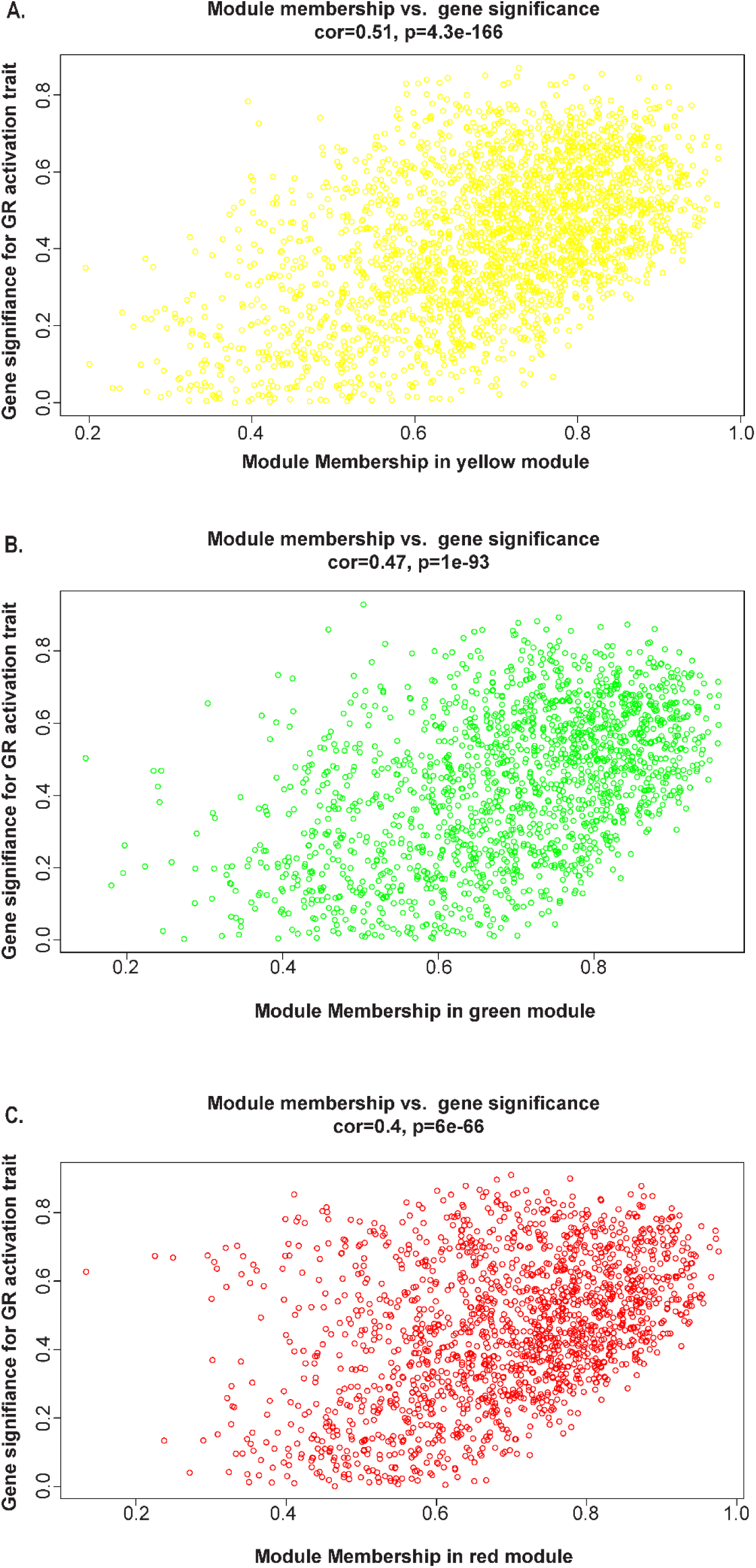
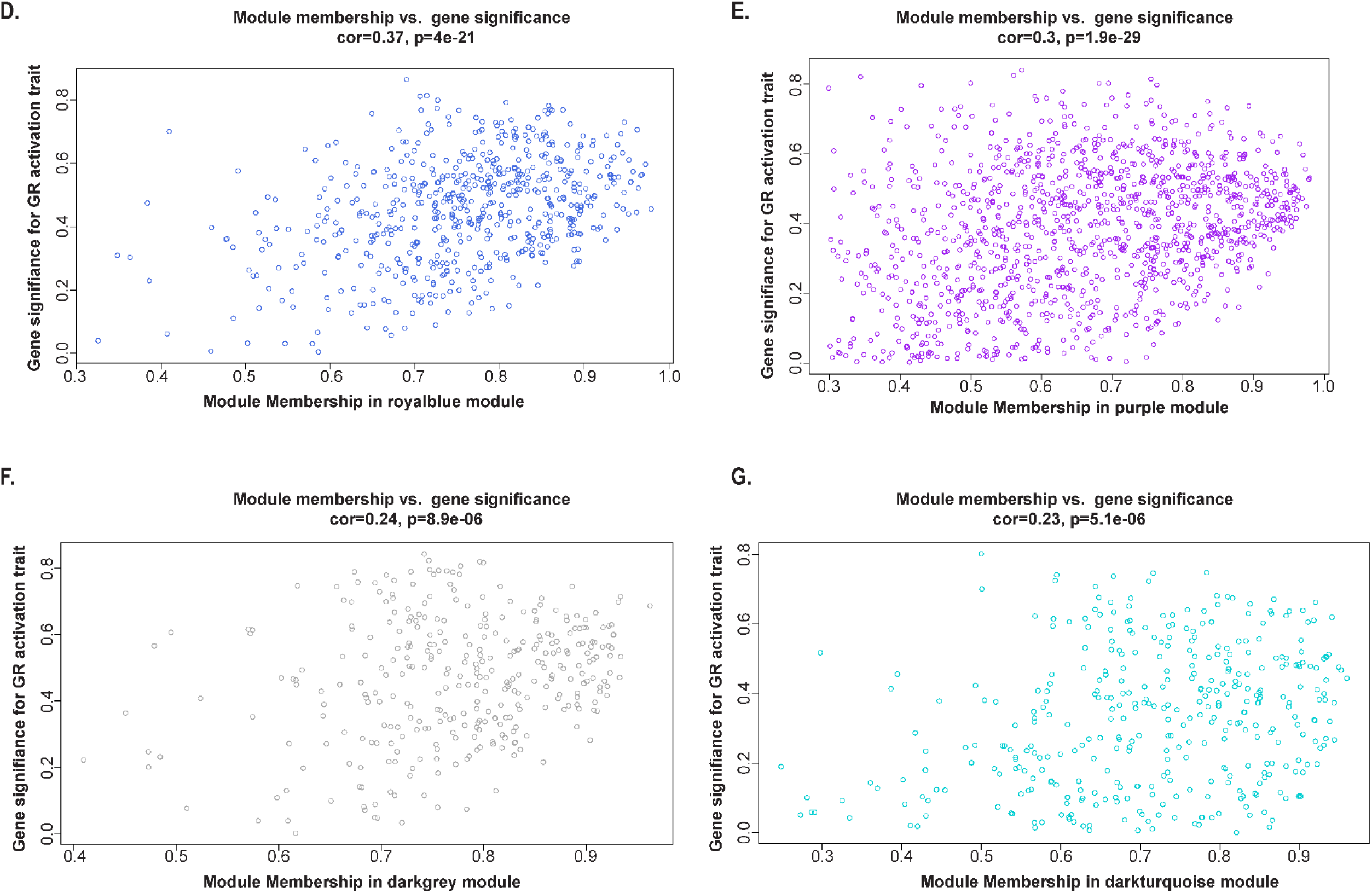
Module building. To generate network using WGCNA, proteins were clustered based on their correlation matrix. This was performed using RNA-Seq data from all 15 samples included in the present study. **Supplementary figure 1 ii-iii.** **A-G.** Correlation between MM and GS of all genes in each module. The strength of correlation is indicated as cor, and p value of the correlation is also indicated for each module.

**Supplementary figure 2.**
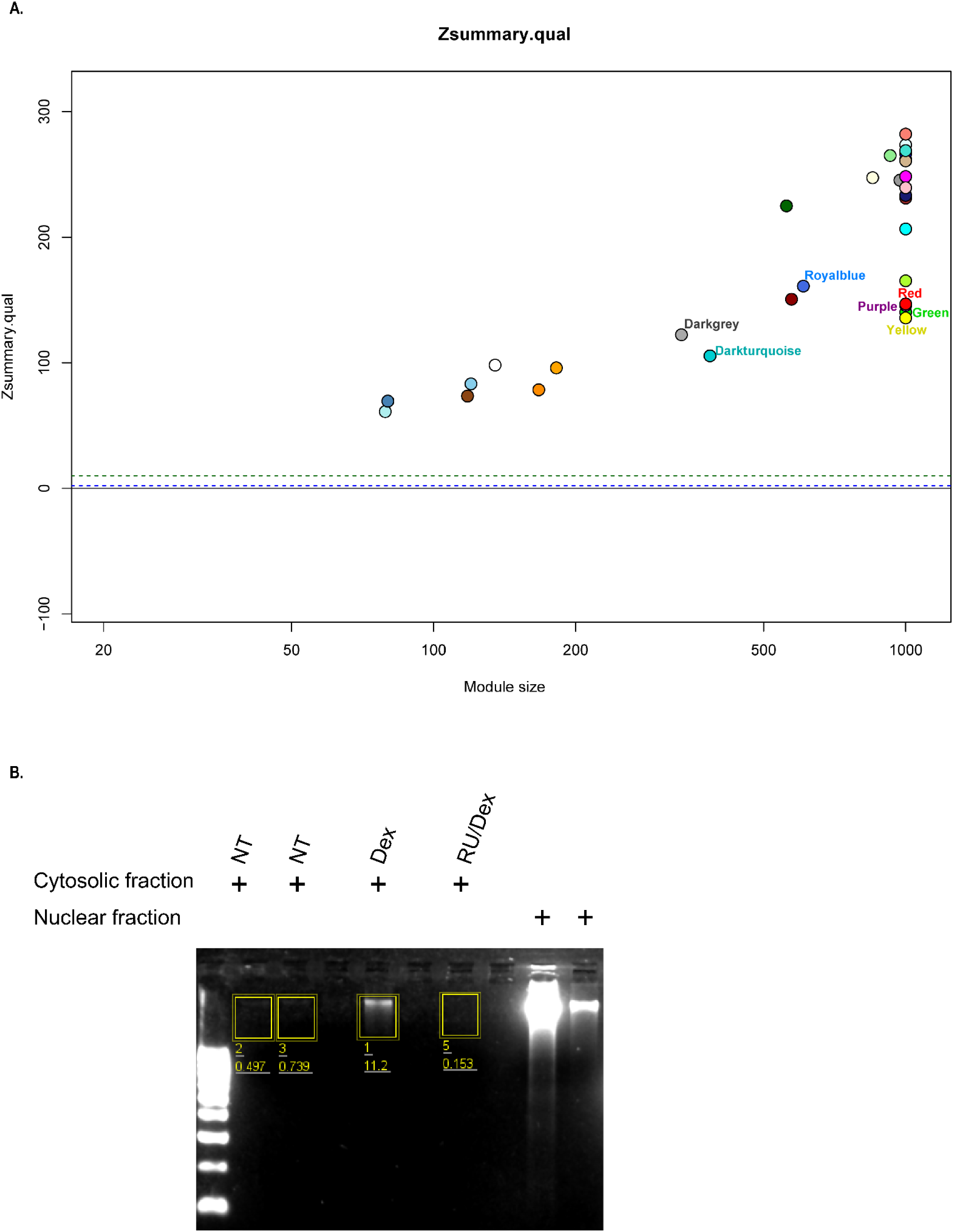
**A.** Zsummary Module quality score (how well-defined modules are in repeated random splits of the reference dataset). Permutation tests were performed to adjust the quality statistics of each module for random chance by defining Z statistics. All modules demonstrated strong evidence for high quality (Zsummary > 10), confirming that the modules identified in the reference network were well-defined and non-random. **B.** Agarose gel analysis of cytosolic extracts from NT iPS-Mg, Dex pre-treated iPS-Mg and RU/Dex pre-treated iPS-Mg.

**Supplementary figure 3.**
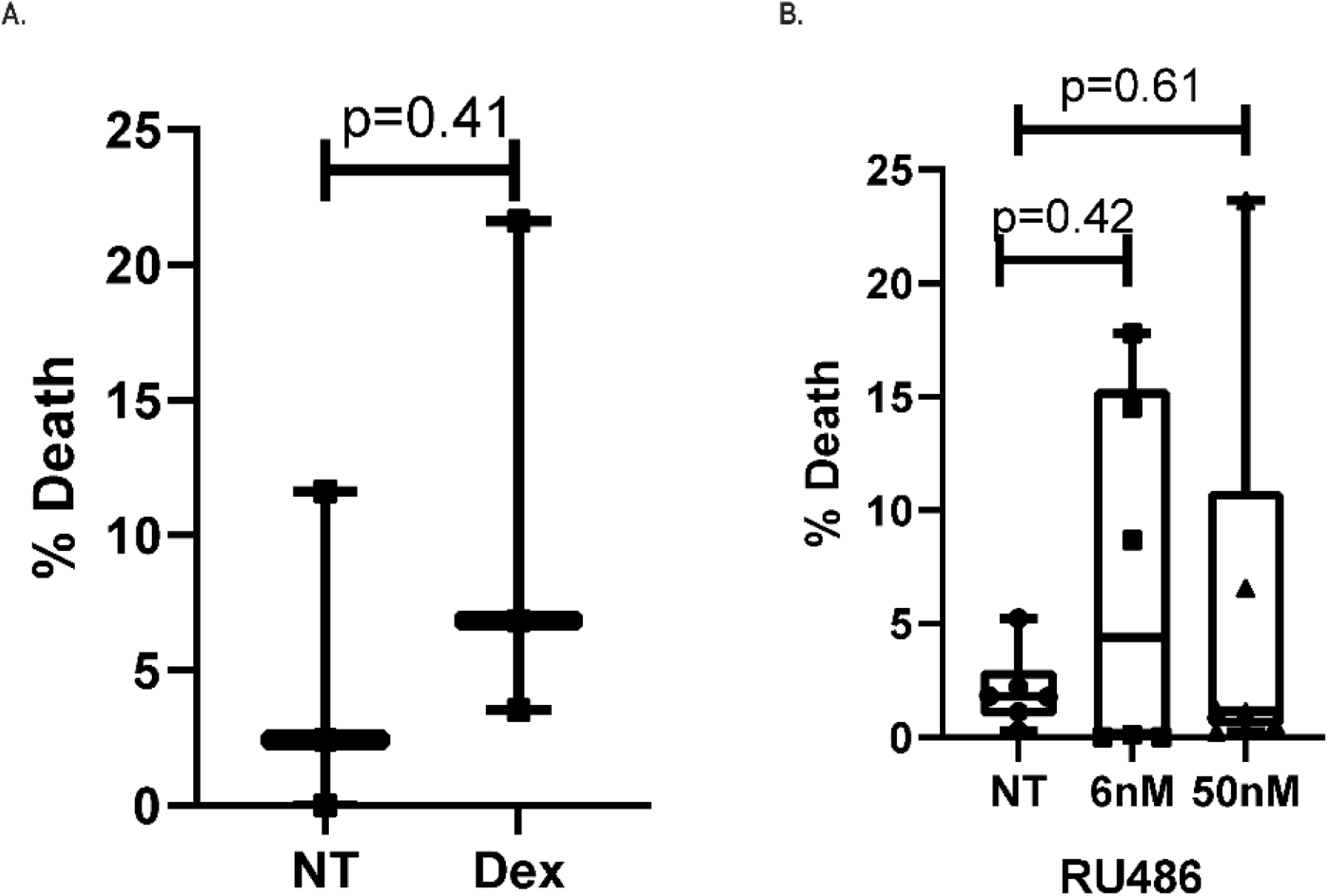
**A.** Quantification of PI staining to determine the % of dead cells in NT, Dex, RU/Dex pretreated iPS-Mg. N=3 independent experiments. Unpaired *t*-tests was used. Box-and-whisker plot representing all data points and the median. **B.** Quantification of PI staining to determine the % of dead cells in NT, 6nM RU486, 50nM RU486 treated iPS-MPro for 24 hours. N=2 independent experiments. Ordinary one-way ANOVA test was used. Box-and-whisker plot representing all data points and the median.

## Materials and Methods

### iPSC-derived microglia and treatment

This study was covered by ethics reference 09/H0716/64, from the National Hospital for Neurology and Neurosurgery and the Institute of Neurology’s joint research ethics committee. A material transfer agreement between University College London and the University of California Irvine Alzheimer’s Disease Research Center enabled the acquisition of R47H heterozygous (R47H^het^) patient-derived fibroblasts (UCI ADRC; Professor M Blurton-Jones) which we previously derived into iPSCs^19^ (Piers et al. 2020). The following lines and clones were used 26. 5, 26. 15, and 8. 6 of the R47H^het^ iPSC. In addition, the following lines from normal individuals expressing common variant TREM2 used in this study were SFC840 (Stembancc), BIONi010-C (EBiSC) and KOLF2_C1 (kindly provided by the Sanger Institute).

Using our previously described protocol, iPS-microglia (iPS-Mg) were generated^19,20,21,22^ (Garcia-Reitboeck et al. 2018; Xiang et al. 2018; Piers et al. 2020, Cosker et al. 2021). All iPSCs were cultured and passaged routinely in Essential 8 medium (Gibco), iPS-myeloid progenitor cells (iPS-MPro) were harvested from the flask of embryoid bodies and further differentiated with IL-34, MCSF and TGF-β1 for 14 days, plus a final maturation with CX3CL1 and CD200 (all growth factors were from Peprotech, London UK) for 3 more days as previously described^19, 23^ (Piers et al. 2020; Mallach et al. 2021).

To model ELS, 50 nM Dex (Tocris Bioscience, Abingdon, UK) or 500 nM hydrocortisone (Cort; Tocris Bioscience, Abingdon, UK) was added during the second stage of primitive haematopoiesis, which typically occur 2 weeks after the stem cell plating, when the embryoid bodies were first seeded into flasks, and subsequently at every week for approximately 12 weeks thereafter (Figure 1A.) when the medium was replenished. These cells are referred to as iPS-myeloid progenitors (iPS-MPro). To block GR activation, 6 nM RU486 (Cayman, CAS#84371-65-3, USA) was added 30 min before 50 nM Dex. This GR antagonistic paradigm has been used other *in vitro* studies^24, 25^ (Liu et al. 2017, Guess et al. 2010). The concentration of 6 nM of RU486 is based on the IC_50_ of RU486^26^ (Attardi et al. 2004). Thus, throughout the rest of this paper, Dex-iPS-Mg means that at the iPS-MPro stage, the cells were treated with Dex, and Cort-iPS-Mg means that at the iPS-MPro stage, cells were treated with Cort, and RU/Dex iPS-Mg (RU486 and Dex treated iPS-Mg) means that at the iPS-MPro stage, cells were treated Ru and Dex. NT-iPS-Mg means the cells at the iPS-MPro stage were not treated.

### Other cell lines

HeLa cells, kindly provided by Dr Jan Attig (The Crick Institute, London UK), were maintained in Dulbecco’s Modified Eagle’s Medium with 10% heat inactivated foetal bovine serum (FBS) (Thermo Fisher Scientific, Gothenberg, Sweden), 10 U/ml Penicillin/Streptomycin (Thermo Fisher Scientific, Dartford, UK), and 1 X GlutaMax (Gibco, Thermo Fisher Scientific). Cells were grown at 6 x 10^6^ density in 100 x 20 mm tissue culture treated petri dish (CytoOne^®^, Starlab, Hamburg, Germany) and maintained at 37°C in a humidified atmosphere with 5% CO_2_. These cells were used as positive controls for glucocorticoid receptor expression, particularly NR3C1.

### Fibroblast culturing

A human primary fibroblast line was maintained in Dulbecco’s Modified Eagle’s Medium with 10% heat inactivated FBS (Thermo Fisher Scientific, Gothenberg, Sweden), 10 U/ml Penicillin/Streptomycin (Thermo Fisher Scientific, Dartford, UK). Cells were grown in a 75 cm^2^ flask (Falcon^®^, Corning Life Science, Flintshire) and maintained at 37°C in a humidified atmosphere with 5% CO_2_. The cells were used with iPSC as positive controls for a relative average telomere assay.

All cell lines were mycoplasma-tested before use according to the manufacturer’s instructions (Mycoalert Detection Kit, Lonza).

### Assessment of Cell Death

Cells were stained for 15 min with Hoechst 33342 (5 μg/ml) to obtain the number of cells present on the coverslip, plus calcein-AM (calcein) to obtain the number of live cells (stained green with calcein-AM, 5 μM) and propidium iodide (PI, 5 μg/ml) to obtain the number of dead cells (staining red) which represent end-state apoptosis or necrotic cells, as previously described^27^ (Davenport et al. 2010). A minimum of 3 fields per coverslip were counted, and at least 3 coverslips were assessed for each variable per experiment.

### Crude cytosolic/nuclear fractionation

Cytoplasmic fractionation was performed to analyse the presence of cytosolic DNA and was carried out according to Song et al. ^28^(2019), with minor modifications. Briefly, 1 million cells were harvested and lysed for 5 min on ice in 250 µl of cytoplasmic fraction buffer (10 mM HEPES, 10 mM KCl, 1. 5 mM MgCl_2_, 0. 34 M sucrose, 10% (v/v) glycerol, pH 7. 0-7. 6 plus Halt™ protease and phosphatase inhibitor cocktail (Thermo Fisher Scientific, Dartford, UK). Subsequently, the pelleted nuclei were removed by low-speed centrifugation (1500 g, 10 min, 4°C). Then 200 µl of the cytoplasmic fraction was processed for genomic DNA extraction using the commercial PureLink^™^ genomic DNA Mini kit (Invitrogen, Thermo Fisher Scientific) according to the manufacturer’s instructions. 10 µl of the cytoplasmic fraction was analysed by BCA assay to normalise cytoplasmic DNA loading to cellular protein. Subsequently, an appropriate volume of the nuclear fraction to provide a positive DNA signal was suspended in 200 µl of PBS and genomic DNA extraction performed. Finally, DNA gel loading dye (Thermo Scientific™, Dartford, UK) was added into the cytoplasmic and nuclear extracts to stain the genomic DNA, and the cytoplasmic DNA concentrations were adjusted according to the protein concentration of cytoplasmic fractions and analysed on a 1-2% agarose gel.

### PCR and quantitative PCR

Cell were plated at 5 x10^5^ per well on a 6-well culture plate. RNA was extracted using RNeasy Plus Minikits (Qiagen, Manchester, UK) and reverse transcribed (RT) with a high-capacity cDNA Reverse Transcription kit (Applied Biosystems, Thermo Fisher). Commercially purchased human primary microglia cDNA (ScienCell, Carlsbad, USA) was also analysed as a control sample.

For PCR analysis of GR-α and β, the PCR master mix (New England Biolab, Hitchin, UK) was prepared according to the manufacturer’s instruction. One µl of cDNA from each sample was used for the amplification of the specific sequences of the GR spice variant. The primers used for GRα and GRβ were purchased from Integrated DNA Technologies (IDT, Belgium), GR total from SelectScience^®^ (Merck, USA), are listed in Table 3.

**Table 3.**
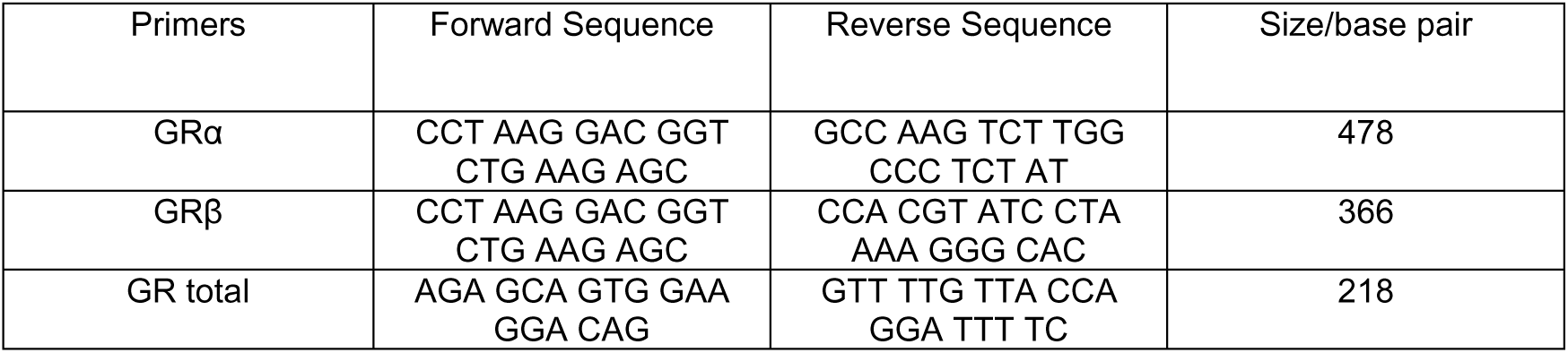
- Primer information for GR analysis

For analysing TMEM119 expression (Hs01938722_u1, Applied Biosystems™), qPCR was carried out using TaqMan Universal Mastermix (4440038, Life Technologies,) in the Stratagene Mx3000p qPCR system and MxPro qPCR software. Expression was normalised to GAPDH (Hs02758991_g1, Applied Biosystems™). For analyzing GR total expression, qPCR was performed using qPCR *Power* SYBR™ Green PCR Master Mix (4367659, Applied Biosystems™) and GR total primer listed in Table 3, and a QuantStudio™ 7 Flex Real-Time PCR system (Applied Biosystems™).

### Relative telomere length measurement using qPCR

Using the crude cytosolic/nuclear fractionation, the nuclear fraction was further purified with 200 µl of PBS (with MgCl_2_ and CaCl2) for 5 min at 900 g to pellet the nuclear fraction further. This is to minimise confounding signals from the micronuclei present in the cytosolic fraction, as micronuclei contain a telomere signal^29^ (Norppa and Falck 2003). The purified nuclear fraction was then used for the telomere assay.

Average telomere measurements using qPCR have previously been described^30, 31^ (Cawthon 2002, Joglekar et al. 2020). We followed the same protocol as Joglekar et al. ^31^ (2020) and chose the single-copy gene (36B4) as a reference based on its primer specificity and efficiency (Table 4). After thermal cycling was completed, Agilent Aria 1. 8 software was used to generate Ct values for telomere signals and reference gene signals. Average Ct value from three technical replicate was input for evaluation of relative telomere length as previously described^32^ (Vasilishina et al. 2019). Telomere to Single copy gene ratio (T/S ratio) was first calculated, then normalised to Ct value of reference DNA sample. Old-age human primary fibroblasts and iPSC were used as positive controls for the telomere length assay. Ct values of telomere and reference signals from iPSC served as a reference DNA sample.

**Table 4.**
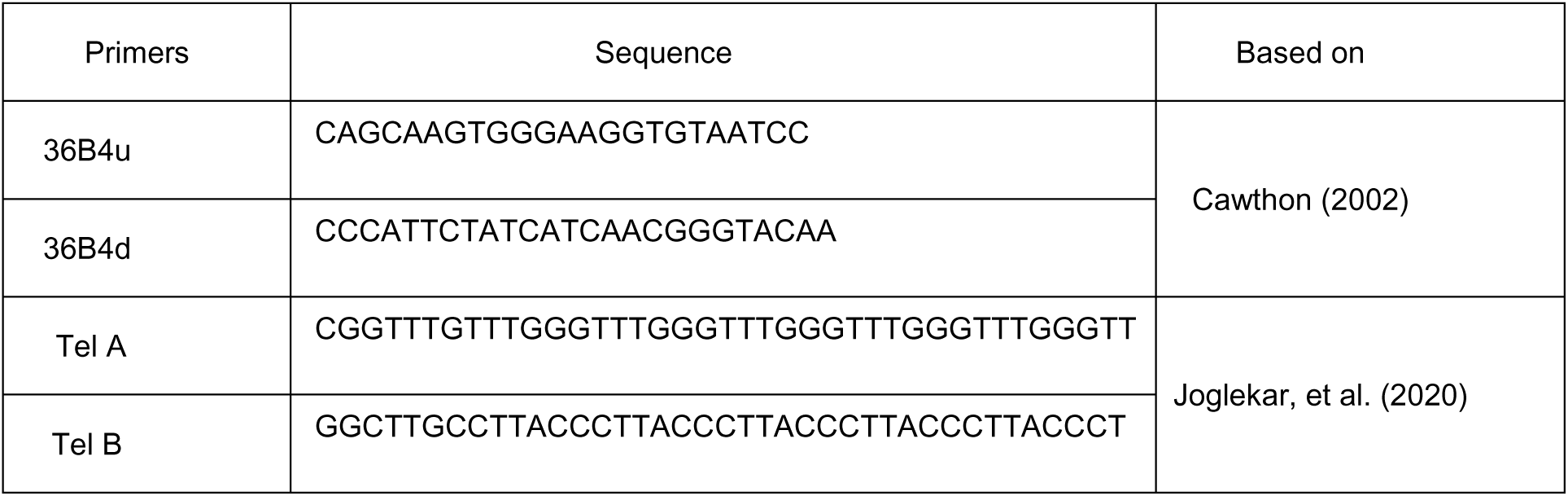
Primers used for Telomere qPCR assay

### Immunocytochemistry of micronuclei

Cells were typically plated at a density of 2 x 10^4^ on glass coverslips in 24 well plates. To stain for micro-nuclei, two antibodies were used, lamin a/c and cGAS. Cells was costained with antibody to P2Y12 to delineate the cytoplasmic space. iPS-Mg were fixed in ice-cold PBS with 4% paraformaldehyde (PFA) for 15 min. The cells were quenched in 50 mM NH_4_Cl for 10 min, followed by permeabilisation with 0. 2% Triton X-100 for 5 min. Primary antibody to lamin a/c (Abcam, Cambridge, UK) at 1:200 dilution was incubated overnight at 4°C in 5% normal goat serum followed by an appropriate secondary fluorescence-conjugated antibody. To stain for cGAS, cells were fixed in ice cold 100% methanol at -20 °C for 5-7 min followed by 2 min brief permeabilization with 0. 2% Triton X-100 in PBS. Primary antibody to cGAS (Santa Cruz Biotechnology, Texas, USA) and P2RY12 (ATLAS, Stockholm, Sweden) at 1:200 dilution was incubated overnight at 4°C in 5% normal goat serum followed by an appropriate secondary fluorescence-conjugated antibody.

Secondary antibodies used were, goat anti-mouse-488 (Life Technologies, Thermo Fisher), or goat anti-rabbit-568 (Life Technologies, Thermo Fisher) at 1:500 dilution for 1 h at RT and nuclei and micronuclei were counterstained with DAPI (1 μg/ml of 4′,6-diamidino-2-phenylindole dihydrochloride in Vectashield Antifade Mounting Medium with DAPI). All images were acquired on a Zeiss LSM710 confocal microscope using the LSM Pascal 5. 0 software. One hundred 63 x magnification tiles from 5-8 random fields of each coverslip were captured. Images were analysed in ZEN 3. 4 (blue edition) software, and co-localisation of lamin a/c with micronuclei or co-localisation of cGAS with micronuclei were manually counted. More than 1500 cells from each condition were counted for lamin a/c and micronuclei; and over 4700 cells from each condition were counted for cGAS with micronuclei. Only non-apoptotic and non-necrotic cells were analysed. Micronuclei classification was defined by criteria based on Ye et al. ^33^ (2019).

### Immunoblotting

Immunoblotting was performed to determine the expression of lamin a/c and GAPDH. Typically, 20 - 40 µl of cytosolic and 60 µl nuclear fraction (isolated as above) were denatured and resolved by SDS-PAGE, transferred onto nitrocellulose membranes and incubated with primary and secondary antibodies (Table 5). To eliminate the possible interference with the protein, presence of DNA in the cytosolic fraction was eliminated using 10µl of DNase I (Thermo Fisher Scientific, Dartford, UK.). Proteins were visualized with an Odyssey detection system (LI-COR) and quantified using ImageJ software (www.imagej.nih.gov/ij).

**Table 5.** Antibodies used for Western blotting

| Target | Cat No | Supplier | Dilution |
| --- | --- | --- | --- |
| GAPDH | AM4300 | Invitrogen | 1:1000 |
| Lamin a/c | Ab238303 | Abcam | 1:1000 |
| Goat Anti-mouse 680 | A21058 | Invitrogen | 1:1000 |
| Goat Anti-mouse 790 | A11357 | Invitrogen | 1:1000 |

### Senescence-associated β galactosidase staining

Senescence was determined with a Senescence Detection Kit (BioVision, Mountain View, CA, USA), according to the manufacturer’s instructions to detect senescence associated β-galactosidase activity (SA-β-gal). A minimum of 5-7 fields of view, in 2 biological replicates, repeated in 4 independent experiments were collected using Zeiss Axiophot light microscope (Carl Zeiss, Oberkochen, Germany) and photographed with a Nikon D300 camera (Nikon Instruments, Melville, NY), equipped with camera control pro 2 software. All images were converted to grey-scale and the same intensity threshold level was set for each image from each experiment. Mean β-gal activity in each field was calculated by using the mean grey value per field divided by the cell number counted per field.

### Senescence associated secretory phenotype

IPS-Mg medium supernatants from 2-3 independent experiments were collected, centrifuged at 300 g for 10 min at 4°C, to remove debris and processed according to the manufacturers’ instructions (Proteome Profiler™ Human XL cytokine array kit; R&D Systems, Abingdon, UK). In total, supernatants pooled from 2-3 biological replicates from NT, Dex, or RU/Dex pre-treated microglia were incubated per membrane, for a total of 2 membranes per condition. Each membrane was imaged with an Odyssey FC (Li-Cor Biosciences, US), and dot density quantified in Image studio Lite Version 5. 2, normalising to both reference and negative control dots.

### Ki-67 proliferation assay

One x 10^6^ iPS-MPro were collected from the flasks and plated on 100 x 20 mm tissue culture treated petri dishes for 72 h to proliferate over a large surface area. Cells were fixed using 80% ethanol and left at -20°C for 1 h before staining. Cells were stained with FITC-conjugated anti-Ki-67 antibody (Miltenyi Biotec, Germany) as per manufacturer’s protocol. To rule out unspecific binding of the Ki-67 antibody, an isotype control antibody was used (1:50 dilution, REA control antibody, Miltenyi Biotec, Germany). Samples were analysed using a flow cytometer (Becton Dickinson FACSCalibur). At least 10 000 events were acquired per condition) and data analysed with Flowing software 2. 5. 1, (University of Turku, Finland).

### Bulk cells RNA-Seq

Total cellular RNA was extracted using the RNeasy Plus Mini Kit (Qiagen, Manchester, UK) according to the manufacturer’s instruction with minor modifications. Multiple replicates from each cell line and each condition were pooled together for RNA extraction and RNA from all samples was submitted to UCL Genomics processing (Zayed Centre for Research into Rare Disease in Children, 20 Guilford Street, London WC1N 1DZ). The concentration and quality (integrity) of the total RNA was determined using a TapeStation System (Agilent) and the RIN (RNA Integrity Number) Quality Metric. RIN values of all samples sent for RNA-Seq were above 7. Thus, 500 ng of RNA were enriched for polyA tailed mRNA, fragmented, reverse transcribed and ligated to index sequencing adapters using Kapa mRNA HyperPrep kit (Roche) according to the manufacturer’s instructions. Amplified libraries were then sequenced on a NovaSeq 6000 SP flow-cell generating 20-25 million reads with 100 bp single-end reads.

### Analysis of RNA-Seq

Raw RNA-seq reads were processed for quality check and adapter trimming using fastp, then aligned to the ensemble human reference GRCh38 genome using HISAT2 (version 2. 2. 1)^34^ (Kim et al. 2019). Gene expression was subsequently quantified using the featureCounts function within Rsubread (version 1. 34. 7)^35, 36^ (Liao et al. 2014, Liao et al. 2019). Then, the raw gene counts were processed with the DESeq2 package (version 1. 26. 0)^37^ (Love, et al. 2014) in R (version 3. 6. 1) to analyse differentially expressed genes (DEGs). Gene Ontology enrichment analysis of DEGs was performed in R using the package clusterProfiler^38^ (version 3. 14. 3, qvalueCutoff = 0. 05) (Yu, et al. 2012). GSEA analysis of the transcriptome was also performed in R using the package clusterProfiler.

### Transcriptomic comparative analysis of in-house iPS-Mg and human primary microglia

Raw FASTQ files of *ex vivo* human primary microglia from different ages (0. 42 – 90 years old)^39,40,41,42^ (Zhang et al. 2016, Galatro et al. 2017, Gosselin et al. 2017, Olah et al. 2018) were downloaded and input for downstream analysis using the same pipeline as our in-house iPS-Mg RNA-Seq data. Briefly, FASTQ files were processed for quality check and adapter trimming using fastp, then pseudo-mapped to the human transcriptome (human Gencode release 36) using Salmon (version 1. 4. 0). Tximport was then used to import Salmon’s transcript-level quantifications and aggregate them to the gene level quantification. The size and depth of the sequencing library per sample and the length of the genes were then normalised using the normalization Factors function within DESeq2. Similarity and distance among samples from different ages of primary microglia and in-house iPS-Mg were assessed using the R default dist function. Before performing sample distances, counts underwent variance stabilizing transformation.

### Comparative Pathway Enrichment Analysis of *in vitro* and *in vivo* model of ELS

To identify pathway enrichments, differentially expressed genes from in house iPSC-Mg treated with Cort were submitted for PANTHER overrepresentation analysis. A comparable analysis was performed on *in vivo* ELS data^43^ (Delpech et al 2016).

### Weighted Gene Co-expression Network Analysis

The featureCount processed genes matrix from the RNA-Seq analysis was first filtered for missing values and zero-variance-genes using the goodSamplesGenes function, and then 36599 genes were included in the WGCNA analysis. One signed network was constructed using the WGCNA package in the R language^44^ (Langfelder and Horvath 2008), with GR activation and genotype as traits. Thus, we performed the analysis of network topology for various soft-thresholding powers to have relative balanced scale independence and mean connectivity of the WGCNA. Finally, the network construction and module detection followed these default parameters: Pearson’s correlations, power of 12, maxBlockSize of 100000, minModuleSize of 10, deepSplit of 2, and mergeCutHeight of 0. 25.

To ensure that the modules identified were indeed associated with the traits, we first determine module significance (MS), which can be defined as the average gene significance (GS) across module genes. GS is a term that refers to the relationship between an individual gene and a biological trait. Thus, modules with a higher mean GS indicate a greater degree of association between the module’s genes and the traits. For the purpose of selecting gene modules of interest, we compared the average GS of the modules. Second, we created a scatter plot of the module membership (MM) versus the GS. Module membership (MM) is defined by WGCNA as *MM*(*i*) = *cor*(*x _i_*, *ME*), and it is used to quantify a gene’s importance within a module.

This is to show that within each module, the genes that are associated with the trait are also the inevitable components of the modules. For instance, if GS and MM were highly correlated, this indicated that genes were critical components of modules and were highly correlated with the trait^44^ (Langfelder and Horvath 2008). The Zsummary. qual module quality and robustness metric was developed by performing a module preservation analysis on a single network as a test and reference^45^ (Langfelder et al. 2011). The modules were functionally annotated using the Gene Ontology enrichment analysis function in R, which is included in the package clusterProfiler (version 3. 14. 3, qvalueCutoff = 0. 05).

### Statistical analysis

Statistical analysis was carried out in GraphPad Prism using unpaired t-test, Mann-Whitney U test, ANOVAs (one-way) and Krushal-Wallis test with Dunn’s multiple comparison tests from at least 3 independent experiments. Statistical significance was defined as *ρ<0. 05, **ρ<0. 01, ***ρ<0. 005. The application of the statistical analysis test was pre-determined by normality testing conducted in Prism.

### Data availability

Accession codes: Gene Expression Omnibus GSE73721^39^ (Zhang et al. 2016), GSE99074^40^ (Galatro et al. 2017), dbGaP: phs001373. v2. p2^41^ (Gosselin et al. 2017), SYNAPSE: syn3219045, syn11468526^42^ (Olah et al. 2018).

## Abbreviations

ADHD: attention-deficit hyperactivity disorder
ASD: autistic spectrum disorder
cGAS: Cyclic GMP-AMP synthase
Cort: hydrocortisone
Dex: dexamethasone
cor: correlation
DEG: differentially expressed gene
ELS: early life stress
FBS: foetal bovine serum
GC: glucocorticoids
GR: glucocorticoid receptors
GS: gene significance
GSEA: Gene set enrichment analysis
h: hr, hours
hMg: human microglia
IFN: interferon
iPS-Mg: human induced pluripotent stem cell derived microglia
iPS-MPro: iPSC-derived myeloid progenitor
ISG: interferon-stimulated genes
MR: mineralocorticoid receptor
MM: module membership
MS: module significance
NT: non-treated
RU: RU486 GR antagonist
SA-β-gal: senescence-associated beta galactosidase
SASP: senescence-associated secretory phenotype
TOM: topological overlap matrix
TREM2: triggering receptor expressed on myeloid cells 2
WGCNA: weighted gene co-expression network analysis

## References

1. Lupien, S. J., B. S. McEwen, M. R. Gunnar and C. Heim (2009). Effects of stress throughout the lifespan on the brain, behavour and cognition. Nature Review Neuroscience 10(6): 434–445.

2. Manzari, N., K. Matvienko-Sikar, F. Baldoni, G. W. O’Keeffe and A. S. Khashan (2019). Prenatal maternal stress and risk of neurodevelopmental disorders in the offspring: a systematic review and meta-analysis. Social psychiatry and psychiatric epidemiology 54(11): 1299–1309.

3. Krontira, A. C., C. Cruceanu and E. B. Binder (2020). Glucocorticoids as Mediators of Adverse Outcomes of Prenatal Stress. Trends Neurosci 43(6): 394–405.

4. Benediktsson, R., R. S. Lindsay, J. R. Seckl and C. R. W. Edwards (1993). Glucocorticoid exposure in utero: new model for adult hypertension. The Lancet 341(8841): 339–341.

5. Swanson, A. M. and A. L. David (2015). Animal models of fetal growth restriction: considerations for translational medicine. Placenta 36(6): 623–630.

6. Quinn, M.A., A. McCalla, B. He, X. Xu and J.A. Cidlowski (2019). Silencing of maternal hepatic glucocorticoid receptor is essential for normal fetal development in mice. Communications Biology 2(1): 104.

7. Chen, W., N. Liu, S. Shen, W. Zhu, J. Qiao, S. Chang, J. Dong, M. Bai, L. Ma, S. Wang, W. Jia, X. Guo, A. Li, J. Xi, C Jiang and J. Kang (2021). Fetal growth restriction impairs hippocampal neurogenesis and cognition via Tet1 in offspring. Cell Reports 37(5): 109912.

8. Schafer, D. P., E. K. Lehrman, A. G. Kautzman, R. Koyama, A. R. Mardinly, R. Yamasaki, R. M. Ransohoff, M. E. Greenberg, B. A. Barres and B. Stevens (2012). Microglia sculpt postnatal neural circuits in an activity and complement-dependent manner. Neuron 74(4): 691–705.

9. Cunningham, C. L., V. Martínez-Cerdeño and S. C. Noctor (2013). Microglia regulate the number of neural precursor cells in the developing cerebral cortex. J Neurosci 33(10): 4216–4233.

10. Squarzoni, P., G. Oller, G. Hoeffel, L. Pont-Lezica, P. Rostaing, D. Low, A. Bessis, F. Ginhoux and S. Garel (2014). Microglia modulate wiring of the embryonic forebrain. Cell Rep 8(5): 1271–1279.

11. Weinhard, L., G. di Bartolomei, G. Bolasco, P. Machado, N. L. Schieber, U. Neniskyte, M. Exiga, A. Vadisiute, A. Raggioli, A. Schertel, Y. Schwab and C. T. Gross (2018). Microglia remodel synapses by presynaptic trogocytosis and spine head filopodia induction. Nature Communications 9(1): 1228.

12. Mondelli, V., A. C. Vernon, F. Turkheimer, P. Dazzan and C. M. Pariante (2017). Brain microglia in psychiatric disorders. Lancet Psychiatry 4(7): 563–572.

13. Courchesne, E., V. H. Gazestani and N. E. Lewis (2020). Prenatal Origins of ASD: The When, What, and How of ASD Development. Trends Neurosci 43(5): 326–342.

14. Monier, A., H. Adle-Biassette, A.-L. Delezoide, P. Evrad. P. Gressens and C. Verney (2007). Entry and distribution of microglial cells in human embryonic and foetal cerebral cortex. Journal of Neuropathology & Experimental Neurology 66(5): 372–382.

15. Ginhoux, F., M. Greter, M. Leboeuf, S. Nandi, P. See, S. Gokhan, M. F. Mehler, S. J. Conway, L. G. Ng, E. R. Stanley, I. M. Samokhvalov and M. Merad (2010). Fate Mapping Analysis Reveals That Adult Microglia Derive from Primitive Macrophages. Science 330(6005): 841–845.

16. Bittle, J. and H. E. Stevens (2018). The role of glucocorticoid, interleukin-1β, and antioxidants in prenatal stress effects on embryonic microglia. J Neuroinflammation 15(1): 44.

17. Park M. J., H. S. Park, M. J. You, J. Yoo, S. H. Kim and M. S. Kwon (2019). Dexamethasone Induces a Specific Form of Ramified Dysfunctional Microglia. Mol Neurobiol. 56(2):1421–1436.

18. Miller, K. N., S. G. Victorelli, H. Salmonowicz, N. Dasgupta, T. Liu, J. F. Passos and P. D. Adams (2021). Cytoplasmic DNA: sources, sensing, and role in aging and disease. Cell 184(22): 5506–5526.

19. Piers TM, Cosker K, Mallach A, Thomas Johnson G, Guerreiro R, Hardy J, Pocock JM. (2020) A locked immunometabolic switch underlies TREM2 R47H loss of function in human iPSC-derived microglia. FASEB J 2020 Feb;34(2):2436–2450. doi: 10.1096/fj.201902447R. Epub 2019 Dec 23. PMID: 31907987.

20. Garcia-Reitboeck P, Phillips A, Piers TM, Villegas-Llerena C, Butler M, Mallach A, Rodrigues C, Arber CE, Heslegrave A, Zetterberg H, Neumann H, Neame S, Houlden H, Hardy J, Pocock JM. (2018) Human Induced Pluripotent Stem Cell-Derived Microglia-Like Cells Harboring TREM2 Missense Mutations Show Specific Deficits in Phagocytosis. Cell Rep. 24(9):2300–2311. doi:10.1016/j.celrep.2018.07.094. PMID:30157425

21. Xiang X, Piers TM, Wefers B, Zhu K, Mallach A, Brunner B, Kleinberger G, Song W, Colonna M, Wurst W, Pocock JM, Haass C (2018) The Trem2 R47H Alzheimer’s risk variant impairs splicing and reduces Trem2 mRNA and protein in mice but not in humans. Molecular Neurodegeneration 13:49–63

22. Cosker K, Mallach A, Limaye J, Piers TM, Staddon J, Neame SJ, Hardy J, Pocock JM (2021) Microglial signalling pathway deficits associated with the patient derived R47H TREM2 variants linked to AD indicate inability to activate inflammasome. Scientific Reports Sci Rep. 2021; 11: 13316. Published online 2021 Jun 25. doi: 10.1038/s41598-021-91207-1.

23. Mallach A, Gobom J, Arber C, Piers TM, Hardy J, Wray S, Zetterberg H, Pocock JM. Differential stimulation of pluripotent stem cell-derived human microglia leads to exosomal proteomic changes affecting neurons. As part of the Special Issue Microglia in Aging and Neurodegenerative Diseases. Cells 2021, 10, 2866. https://doi.org/10.3390/cells10112866

24. Liu L, Aleksandrowicz E, Schönsiegel F, Gröner D, Bauer N, Nwaeburu CC, Zhao Z, Gladkich J, Hoppe-Tichy T, Yefenof E, Hackert T. Dexamethasone mediates pancreatic cancer progression by glucocorticoid receptor, TGFβ and JNK/AP-1. Cell death & disease. 2017 Oct;8(10):e3064-.

25. Guess A, Agrawal S, Wei CC, Ransom RF, Benndorf R, Smoyer WE. Dose-and time-dependent glucocorticoid receptor signaling in podocytes. American Journal of Physiology-Renal Physiology. 2010 Oct 1.

26. Attardi BJ, Burgenson J, Hild SA, Reel JR. In vitro antiprogestational/antiglucocorticoid activity and progestin and glucocorticoid receptor binding of the putative metabolites and synthetic derivatives of CDB-2914, CDB-4124, and mifepristone. J. Steroid Biochem. Mol. Biol. 2004; 88:277–288.

27. Davenport, C. M., Sevastou, I. G., Hooper, C. and Pocock, J. M. (2010). Inhibiting p53 pathways in microglia attenuates microglial-evoked neurotoxicity following exposure to Alzheimer peptides. Journal of Neurochemistry, 112(2): 552–563. doi: 10.1111/j.1471-4159.2009.06485.

28. Song. X., F. Ma and K. Herrup (2019). Accumulation of Cytoplasmic DNA due to ATM deficiency Activates the Microglial Viral Response System with Neurotoxic Consequences. The Journal of Neuroscience 39(32): 6378–6394.

29. Norppa, H. and G. C.-M. Falck (2003). What do human micronuclei contain? Mutagenesis 18(3): 221–233.

30. Cawthon, R. M. (2002). Telomere measurement by quantitative PCR. Nuclei acids research 30(10): e47–e47.

31. Joglekar, M. V., S. N. Satoor, W. K. M. Wong, F. Cheng, R. C. W. Ma and A. A Hardikar (2020). An optimised step-by-step protocol for measuring relative telomere length. Methods and Protocols 3(2): 27.

32. Vasilishina, A., A. Kropotov, I. Spivak and A. Bernadotte (2019). Relative Human Telomere Length Quantification by Real-Time PCR. Methods Mol Biol 1896: 39–44.

33. Ye, C.J., Sharpe, Z., Alemara, S., Mackenzie, S., Liu, G., Abdallah, B., Horne, S., Regan, S. and Heng, H.H., 2019. Micronuclei and genome chaos: changing the system inheritance. Genes, 10(5), p.366.

34. Kim, D., J. M. Paggi, C. Park, C. Bennett and S. L. Salzberg (2019). Graph-based genome alignment and genotyping with HISAT2 and HISAT-genotype. Nat Biotechnol 37(8): 907–915.

35. Liao, Y., G. K. Smyth and W. Shi (2014). featureCounts: an efficient general purpose program for assigning sequence reads to genomic features. Bioinformatics 30(7): 923–930.

36. Liao, Y., G. K. Smyth and W. Shi (2019). The R package Rsubread is easier, faster, cheaper and better for alignment and quantification of RNA sequencing reads. Nucleic Acids Research 47(8): e47–e47.

37. Love, M. I., W. Huber and S. Anders (2014). Moderated estimation of fold change and dispersion for RNA-seq data with DESeq2. Genome Biol 15(12): 550.

38. Yu, G., L. G. Wang, Y. Han and Q. Y. He (2012). clusterProfiler: an R package for comparing biological themes among gene clusters. Omics 16(5): 284–287.

39. Zhang, Y., S. A. Sloan, L. E. Clarke, C. Caneda, C. A. Plaza, P. D. Blumenthal, H. Vogel, G. K. Steinberg, M. S. Edwards, G. Li, J. A. Duncan, 3rd, S. H. Cheshier, L. M. Shuer, E. F. Chang, G. A. Grant, M. G. Gephart and B. A. Barres (2016). Purification and Characterization of Progenitor and Mature Human Astrocytes Reveals Transcriptional and Functional Differences with Mouse. Neuron 89(1): 37–53.

40. Galatro, T. F., I. R. Holtman, A. M. Lerario, I. D. Vainchtein, N. Brouwer, P. R. Sola, M. M. Veras, T. F. Pereira, R. E. P. Leite, T. Möller, P. D. Wes, M. C. Sogayar, J. D. Laman, W. den Dunnen, C. A. Pasqualucci, S. M. Oba-Shinjo, E. W. G. M. Boddeke, S. K. N. Marie and B. J. L. Eggen (2017). Transcriptomic analysis of purified human cortical microglia reveals age-associated changes. Nature Neuroscience 20(8): 1162–1171.

41. Gosselin, D., D. Skola, N. G. Coufal, I. R. Holtman, J. C. M. Schlachetzki, E. Sajti, B. N. Jaeger, C. O’Connor, C. Fitzpatrick, M. P. Pasillas, M. Pena, A. Adair, D. D. Gonda, M. L. Levy, R. M. Ransohoff, F. H. Gage and C. K. Glass (2017). An environment-dependent transcriptional network specifies human microglia identity. Science 356(6344): eaal3222.

42. Olah, M., E. Patrick, A.-C. Villani, J. Xu, C. C. White, K. J. Ryan, P. Piehowski, A. Kapasi, P. Nejad, M. Cimpean, S. Connor, C. J. Yung, M. Frangieh, A. McHenry, W. Elyaman, V. Petyuk, J. A. Schneider, D. A. Bennett, P. L. De Jager and E. M. Bradshaw (2018). A transcriptomic atlas of aged human microglia. Nature Communications 9(1): 539.

43. Delpech, J. C., L. Wei, J. Hao, X. Yu, C. Madore, O. Butovsky and A. Kaffman (2016). Early life stress perturbs the maturation of microglia in the developing hippocampus. Brain Behav Immun 57: 79–93.

44. Langfelder, P. and S. Horvath (2008). WGCNA: an R package for weighted correlation network analysis. BMC bioinformatics 9(1): 1–13.

45. Langfelder, P., R. Luo, M. C. Oldham and S. Horvath (2011). Is my network module preserved and reproducible? PLoS Comput Biol 7(1): e1001057.

46. Timmermans, S., Souffriau, J. and Libert, C., 2019. A general introduction to glucocorticoid biology. Frontiers in immunology, 10, p.1545.

47. Lieberman, R., H. R. Kranzler, E. S. Levine and J. Covault (2017). Examining FKBP5 mRNA expression in human iPSC-derived neural cells. Psychiatry Res 247: 172–181.

48. Lu, N. Z. and J. A. Cidlowski (2005). Translational regulatory mechanisms generate N-terminal glucocorticoid receptor isoforms with unique transcriptional target genes. Mol Cell 18(3): 331–342.

49. Abud, E. M., R. N. Ramirez, E. S. Martinez, L. M. Healy, C. H. H. Nguyen, S. A. Newman, A. V. Yeromin, V. M. Scarfone, S. E. Marsh, C. Fimbres, C. A. Caraway, G. M. Fote, A. M. Madany, A. Agrawal, R. Kayed, K. H. Gylys, M. D. Cahalan, B. J. Cummings, J. P. Antel, A. Mortazavi, M. J. Carson, W. W. Poon and M. Blurton-Jones (2017). iPSC-Derived Human Microglia-like Cells to Study Neurological Diseases. Neuron 94(2): 278–293.e279.

50. Haenseler, W., S. N. Sansom, J. Buchrieser, S. E. Newey, C. S. Moore, F. J. Nicholls, S. Chintawar, C. Schnell, J. P. Antel, N. D. Allen, M. Z. Cader, R. Wade-Martins, W. S. James and S. A. Cowley (2017). A Highly Efficient Human Pluripotent Stem Cell Microglia Model Displays a Neuronal-Co-culture-Specific Expression Profile and Inflammatory Response. Stem Cell Reports 8(6): 1727–1742.

51. Ozaki, K., D. Kato, A. Ikegami, A. Hashimoto, S. Sugio, Z. Guo, M. Shibushita, T. Tatematsu, K. Haruwaka, A. J. Moorhouse, H. Yamada and H. Wake (2020). Maternal immune activation induces sustained changes in foetal microglia motility. Scientific Reports 10(1): 21378.

52. Binder EB. The role of FKBP5, a co-chaperone of the glucocorticoid receptor in the pathogenesis and therapy of affective and anxiety disorders. Psychoneuroendocrinology. 2009 Dec;34 Suppl 1:S186–95. doi: 10.1016/j.psyneuen.2009.05.021. PMID: 19560279

53. Ke, X., Q. Fu, A. Majnik, S. Cohen, Q. Liu and R. Lane (2018). Adverse early life environment induces anxiety-like behavior and increases expression of FKBP5 mRNA splice variants in mouse brain. Physiological Genomics 50(11): 973–981.

54. Mackenzie, K. J., P. Carroll, C.-A. Martin, O. Murina, A. Fluteau, D. J. Simpson, N. Olova, H. Sutcliffe, J. K. Rainger, A. Leitch, R. T. Osborn, A. P. Wheeler, M. Nowotny, N. Gilbert, T. Chandra, M. A. M. Reijns and A. P. Jackson (2017). cGAS surveillance of micronuclei links genome instability to innate immunity. Nature 548(7668): 461–465.

55. Longui, C. A., M. C. Santos, C. B. Formiga, D. V. Oliveira, M. N. Rocha, C. D. Faria, C. Kochi and O. Monte (2005). Antiproliferative and apoptotic potencies of glucocorticoids: nonconcordance with their antiinflammatory and immunosuppressive properties. Arq Bras Endocrinol Metabol 49(3): 378–383.

56. Singer, G. A. C., A. T. Lloyd, L. B. Huminiecki and K. H. Wolfe (2004). Clusters of Co-expressed Genes in Mammalian Genomes Are Conserved by Natural Selection. Molecular Biology and Evolution 22(3): 767–775.

57. de la Fuente, A. (2010). From differential expression’ to ‘differential networking’ – identification of dysfunctional regulatory networks in diseases. Trends in Genetics 26(7): 326–333.

58. Keating, S. E., M. Baran and A. G. Bowie (2011). Cytosolic DNA sensors regulating type I interferon induction. Trend in Immunology 32(12): 574–581.

59. Honda, K., A. Takaoka and T. Taniguchi (2006). Type I interferon gene induction by interferon regulatory factor family of transcription factors. Immunity 25(3): 349–360.

60. Dubik N, Mai S. Lamin A/C: Function in Normal and Tumor Cells. Cancers (Basel). 2020 Dec 9;12(12):3688. doi: 10.3390/cancers12123688. PMID: 33316938; PMCID: PMC7764147.

61. Kneissig, M., K. Keuper, M. S. de Pagter, M. J. van Roosmalen, J. Martin, H. Otto, V. Passerini, A. Campos Sparr, I. Renkens, F. Kropveld, A. Vasudevan, J. M. Sheltzer, W. P. Kloosterman and Z. Storchova (2019). Micronuclei-based model system reveals functional consequences of chromothripsis in human cells. Elife 8.

62. Kwon M, Leibowitz ML, Lee JH. Small but mighty: the causes and consequences of micronucleus rupture. Exp Mol Med. 2020 Nov;52(11):1777–1786. doi: 10.1038/s12276-020-00529-z. Epub 2020 Nov 24. PMID: 33230251; PMCID: PMC8080619

63. Hatch EM, Fischer AH, Deerinck TJ, Hetzer MW. Catastrophic nuclear envelope collapse in cancer cell micronuclei. Cell. 2013 Jul 3;154(1):47–60. doi: 10.1016/j.cell.2013.06.007. PMID: 23827674; PMCID: PMC3749778.

64. Zierhut, C. and H. Funabiki (2020). Regulation and Consequences of cGAS Activation by Self-DNA. Trends in Cell Biology 30(8): 594–605.

65. Glück S, Ablasser A. Innate immunosensing of DNA in cellular senescence. Curr Opin Immunol. 2019 Feb;56:31–36. doi: 10.1016/j.coi.2018.09.013. Epub 2018 Oct 5. PMID: 30296662.

66. Galvis D, Walsh D, Harries LW, Latorre E, Rankin J. A dynamical systems model for the measurement of cellular senescence. J R Soc Interface. 2019 Oct 31;16(159):20190311. doi: 10.1098/rsif.2019.0311. Epub 2019 Oct 9. PMID: 31594522; PMCID: PMC6833332.

67. Lunyak VV, Armaro-Ortiz A, Gaur Meenakshi (2017) Mesechymal stem cells secretory responses: Senescence messaging secretome and immunomodulation perspective. Fronteirs Genetics. 19 Dec:doi.org/10.3389/fgene.2017.00220.

68. Hagendorf A, Koper J.W, de Jong FH, Brinkmann AO, Lamberts SWJ, Feelders RA, (2005). Expression of the Human Glucocorticoid Receptor Splice Variants α, β, and P in Peripheral Blood Mononuclear Leukocytes in Healthy Controls and in Patients with Hyper- and Hypocortisolism. J. Clin Endocrin Metab, 90(11):m6237–6243, https://doi.org/10.1210/jc.2005-1042.

69. Samuelsson MKR, Pazirandeh A, Davani B, Okret S. (1999) p57^Kip2^, a Glucocorticoid-Induced Inhibitor of Cell Cycle Progression in HeLa Cells. Mol Endocrin 13 (11):1811–1822, https://doi.org/10.1210/mend.13.11.0379

70. Piette C, Deprez M, Thierry Roger T, Noel A, Foidart J-M, and Carine Munau C (2009). The dexamethasone-induced Inhibition of proliferation, migration, and invasion in glioma cell lines is antagonized by macrophage migration inhibitory factor (MIF) and can be enhanced by specific MIF Inhibitors. J Biol Chem 284(47);32483–32493.

71. Cruceanu C, Dony L, Krontira AC, Fischer DS, Roeh S, Di Giaimo R, Kyrousi C, Kaspar L, Arloth J, Czamara D, Gerstner N, Martinelli S, Wehner S, Breen MS, Koedel M, Sauer S, Sportelli V, Rex-Haffner M, Cappello S, Theis FJ, Binder EB. Cell-Type-Specific Impact of Glucocorticoid Receptor Activation on the Developing Brain: A Cerebral Organoid Study. Am J Psychiatry. 2021 Oct 26:appiajp202121010095. doi: 10.1176/appi.ajp.2021.21010095. Epub ahead of print. PMID: 34698522.

72. Zannas AS, Jiaa M, Hafnera K, Baumert J, Wiechmann T, Pape JC, Arloth J, Ködel M, Martinelli S, Roitman M, Röh S, Haehle A, Emeny RT, Iurato S, Carrillo-Roa T, Lahti, J, Räikkönen K, Eriksson JG, Drake AJ, Waldenberger M, Wahl S, Kunze S, Lucae S, Bradley B, Gieger C, Hausch F, Smith AK, Ressler KJ, Müller-Myhsok B, Ladwig, K-H, Rein T, Gassen NC, Binder EB. (2019) Epigenetic upregulation of FKBP5 by aging and stress contributes to NF-κB–driven inflammation andcardiovascular risk. PNAS 116(23): 11370–11379

73. Jameson RR, Seidler FJ, Qiao D, Slotkin TA. Adverse neurodevelopmental effects of dexamethasone modeled in PC12 cells: identifying the critical stages and concentration thresholds for the targeting of cell acquisition, differentiation and viability. Neuropsychopharmacology. 2006 Aug;31(8):1647–58. doi: 10.1038/sj.npp.1300967. Epub 2005 Nov 30. PMID: 16319912.

74. Osathanondh, R., D. Tulchinsky, H. Kamali, M. d. Fencl and H. W. Taeusch (1977). Dexamethasone levels in treated pregnant women and newborn infants. The Journal of Paediatrics 90(4): 617–620.

75. Charles, B., P. Schild, P. Steer, D. Cartwright and T. Donovan (1993). Pharmacokinetics of dexamethasone following single-dose intravenous administration to extremely low birth weight infants. Dev Pharmacol Ther 20(3-4): 205–210.

76. Núñez Estevez, K. J., A. N. Rondón-Ortiz, J. Q. T. Nguyen and A. C. Kentner (2020). Environmental influences on placental programming and offspring outcomes following maternal immune activation. Brain, Behaviour, and Immunity 83: 44–55.

77. Diz-Chaves, Y., M. Astiz, M. J. Bellini and L. M. Garcia-Segura (2013). Prenatal stress increases the expression of proinflammatory cytokines and exacerbates the inflammatory response to LPS in the hippocampal formation of adult male mice. Brain, Behavior, and Immunity 28: 196–206.

78. Diz-Chaves, Y., O. Pernía, P. Carrero, and L. M. Garcia-Segura (2012). Prenatal stress causes alterations in the morphology of microglia and the inflammatory response of the hippocampus of adult female mice. Journal of Neuroinflammation 9(1): 71.

79. Matthews, L. C., A. A. Berry, D. J. Morgan, T. M. Poolman, K. Bauer, F. Kramer, D. G. Spiller, R. V. Richardson, K. E. Chapman, S. N. Farrow, M. R. Norman, A. J. Williamson, A. D. Whetton, S. S. Taylor, J. P. Tuckermann, M. R. White and D. W. Ray (2015). Glucocorticoid receptor regulates accurate chromosome segregation and is associated with malignancy. Proc Natl Acad Sci U S A 112(17): 5479–5484.

80. Flammer, J. R., J. Dobrovolna, M. A. Kennedy, Y. Chinenov, C. K. Glass, L. B. Ivashkiv and I. Rogatsky (2010). The Type I Interferon Signaling Pathway Is a Target for Glucocorticoid Inhibition. Molecular and Cellular Biology 30(19): 4564–4574.

81. Smith, A. M. and M. Dragunow (2014). The human side of microglia. Trends in Neurosciences 37(3): 125–135.

82. Gisselsson, D., J. Björk, M. Höglund, F. Mertens, P. Dal Cin, M. Åkerman and N. Mandahl (2001). Abnormal Nuclear Shape in Solid Tumors Reflects Mitotic Instability. The American Journal of Pathology 158(1): 199–206.

83. Kushima I, Aleksic B, Nakatochi M, Shimamura T, Okada T, Uno Y, Morikawa M, Ishizuka K, Shiino T, Kimura H, Arioka Y. Comparative analyses of copy-number variation in autism spectrum disorder and schizophrenia reveal etiological overlap and biological insights. Cell reports. 2018 Sep 11;24(11):2838–56.

